# OpenDVP: An experimental and computational framework for community-empowered deep visual proteomics

**DOI:** 10.1101/2025.07.13.662099

**Authors:** Jose Nimo, Sonja Fritzsche, Daniela S. Valdes, Minh Trinh, Tancredi Pentimalli, Melissa Klingeberg, Simon Schallenberg, Frederick Klauschen, Florian Herse, Stefan Florian, Nikolaus Rajewsky, Fabian Coscia

**Affiliations:** Max-Delbrück-Center for Molecular Medicine in the Helmholtz Association (MDC), Berlin, Germany; Charité – Universitätsmedizin Berlin, corporate member of Freie Universität Berlin and Humboldt-Universität zu Berlin, Berlin, Germany; Humboldt-Universität zu Berlin, Institute of Biology, Berlin, Germany; Laboratory for Systems Biology of Regulatory Elements, Berlin Institute for Medical Systems Biology (BIMSB), Max-Delbrück-Centrum for Molecular Medicine in the Helmholtz Association (MDC), Berlin, Germany; Institute of Pathology, Charité – Universitätsmedizin Berlin, corporate member of Freie Universität Berlin and Humboldt-Universität zu Berlin, Berlin, Germany; Institute of Pathology, Ludwig Maximilians University Hospital Munich, Munich, Germany; German Cancer Consortium (DKTK), Partner Site Munich, and German Cancer Research Center (DKFZ), Heidelberg, Germany; BIFOLD - Berlin Institute for the Foundations of Learning and Data, Berlin, Germany; Experimental and Clinical Research Center, a cooperation between the Max-Delbrück-Center for Molecular Medicine in the Helmholtz Association and the Charité - Universitätsmedizin Berlin, Germany; Cluster of Excellence, NeuroCure, Charitéplatz 1, 10117 Berlin, Germany; German Cancer Consortium (DKTK), Partner Site Berlin, Heidelberg, Germany; German Center for Cardiovascular Research (DZHK), Site Berlin, Berlin, Germany; National Center for Tumor Diseases (NCT), Site Berlin, Berlin, Germany; German Center for Child and Adolescent Health (DZKJ), Site Berlin, Germany; Max Delbrück Center for Molecular Medicine (MDC) and Cluster of Excellence ImmunoPreCept and Einstein Center for Early Disease Interception (EC-EDI)

## Abstract

Deep visual proteomics (DVP) is an emerging approach for cell type-specific and spatially resolved proteomics. However, its broad adoption has been constrained by the lack of an open-source end-to-end workflow in a community-driven ecosystem. Here, we introduce openDVP, an experimental and computational framework for simplifying and democratizing DVP. OpenDVP integrates open-source software for image analysis, including MCMICRO, QuPath, and Napari, and uses the scverse data formats AnnData and SpatialData for multi-omics integration. It offers two workflows: a fast-track pipeline requiring no image analysis expertise and an artificial intelligence (AI)-powered pipeline with recent algorithms for image pre-processing, segmentation, and spatial analysis. We demonstrate openDVP’s versatility in three archival tissue studies, profiling human placenta, early-stage lung cancer, and locally relapsed breast cancer. In each study, our framework provided insights into health and disease states by integrating spatial single-cell phenotypes with exploratory proteomic data. Finally, we introduce deep proteomic profiling of cellular neighborhoods as a scalable approach to accelerate spatial discovery proteomics across biological systems.

## Introduction

Recent advances in spatial omics have revolutionized our understanding of tissue biology and disease processes ^1^. In particular, spatial proteomics is experiencing remarkable progress in sensitivity, throughput, and spatial resolution ^2^. The ability to map proteins with high spatial resolution has provided novel insights into cellular organization, tissue heterogeneity, and disease mechanisms with unprecedented detail. The significance of these advancements lies in the premise that proteomic profiles provide a direct perspective on the functional and phenotype-centric cellular states that govern health and disease. Historically, spatial proteomics has depended on targeted, antibody-based methodologies ^3^, which offer excellent spatial resolution, yet require prior knowledge for antibody panel design. Antibody-based methods, which generally profile up to ∼60 proteins ^4^, capture a comparatively small fraction of the proteome, which is estimated to encompass more than 10,000 different proteins per single cell type ^5^. A highly synergistic approach involves integrating targeted methods with ultrasensitive liquid chromatography-mass spectrometry (LC-MS)-based proteomics. This multiscale approach has paved the way for exploratory, comprehensive analyses of cell-type and spatially resolved tissue proteomes ^6,7^. Recently, we co-developed deep visual proteomics (DVP)^8^, which combines tissue imaging (immunofluorescence [IF] or immunohistochemistry [IHC]), machine learning-based image analysis, automated laser microdissection (LMD), and ultrasensitive MS for the exploratory proteomic profiling of cell types or regions of interest (ROI). DVP enables the systematic mapping of thousands of proteins, elucidating their networks and signaling pathways within complex tissue architectures, and providing unprecedented insights into health and disease states. For example, DVP recently enabled the discovery of a curative treatment strategy for a fatal human skin disease ^9^ and mapped single-cell proteotoxicity in human liver ^10^.

For image analysis, the Biological Image Analysis Software (BIAS) ^8^ has enabled full support of DVP experiments. However, broader adoption and methodological development have been limited by the absence of non-proprietary, open-source workflows that support scalable image analysis, laser microdissection experimental design, multimodal data integration, and interoperability across imaging and omics platforms. These limitations have constrained systematic benchmarking, extensibility, and community-driven DVP innovations.

To address these gaps, we developed openDVP, an open-source modular framework designed to complement and extend existing DVP solutions, including BIAS. OpenDVP provides flexible, interoperable workflows that support simplified experimental strategies for users with limited computational expertise and advanced pipelines optimized for scalability and complex analyses. Built to integrate state-of-the-art bioimage analysis, spatial statistics, and ultrasensitive mass spectrometry–based proteomics, openDVP enables scalable acquisition and analysis of hundreds of spatially resolved proteomes. Across three archival tissue studies, we demonstrate the versatility of the framework and show that cellular neighborhood–guided proteome profiling offers a balanced alternative to technically demanding single-cell isolation, enabling efficient spatial discovery proteomics at scale.

## Results

### Overview of the openDVP framework

We developed openDVP, an open-source modular framework for user-empowered deep visual proteomics (**Fig. 1a**) (https://github.com/CosciaLab/openDVP). Our end-to-end framework encompasses optimized functions for all the key steps of the DVP workflow, supporting image preprocessing and analysis, automated laser microdissection, interactive image data visualization, and multimodal data integration. OpenDVP supports the most common imaging and proteomics data formats by current and extending functionalities from existing open-source software, such as MCMICRO^11^, SOPA^12^, QuPath ^13^, Napari ^14^, and SpatialData ^15^. Importantly, the modules’ inputs and outputs are standard file formats for seamless integration of community-published or in-house tools. We introduce two complementary workflows: a simplified fast-track pipeline, termed flashDVP, which does not require image analysis expertise, and an extended AI-powered pipeline capable of in-depth spatial tissue analyses. FlashDVP serves as an intuitive entry point for users seeking to rapidly convert raw images (i.e., H&E, IHC, or IF) into exploratory spatial proteomics results. The extended AI-powered DVP workflow enables more experienced users to customize and optimize the pipeline using advanced single-cell spatial analysis tools for more complex phenotype-to-proteotype associations.

**Fig. 1:**
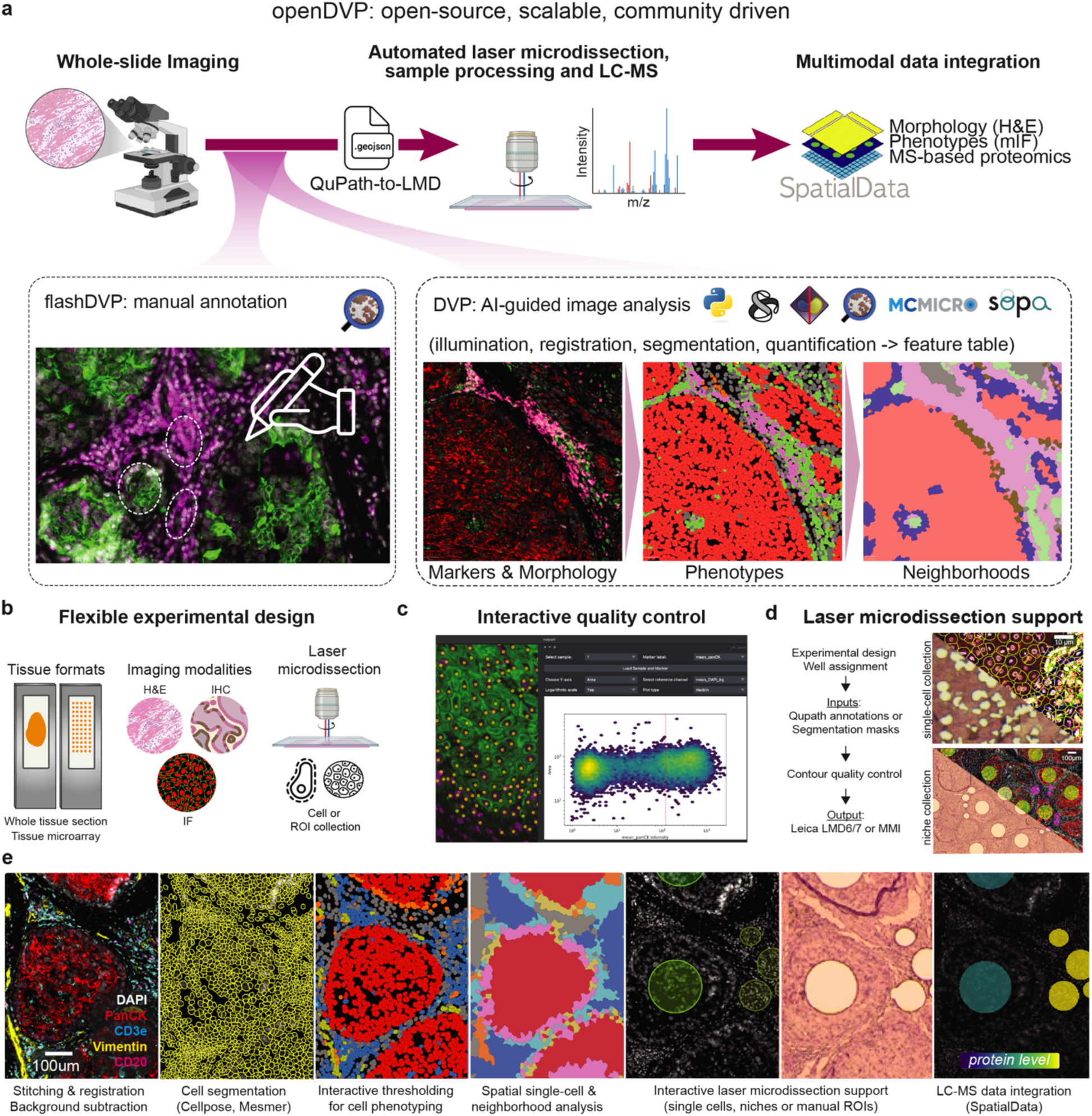
Overview of the openDVP framework. **(a)** Main workflow of openDVP begins with tissue imaging to produce TIFF files. FlashDVP comprises manual image annotation, coordinate transfer with the QuPath-to-LMD webapp, and laser microdissection. OpenDVP provides a framework to process images into feature tables and analyze matrices to phenotype cells, using open-source tools. These data layers provide the basis for contour export and laser microdissection via QuPath-to-LMD. **(b)** OpenDVP supports tissue microarrays or whole-slide images, common imaging modalities, and different LMD collection strategies. **(c)** The Napari-cell-gater plugin enables phenotyping through visual feedback thresholding, plotting cell features in FACS-like density plots. **(d)** QuPath-to-LMD webapp allows users to design, transfer, and validate annotations from QuPath and openDVP to an LMD-ready file format. **(e)** Exemplary openDVP workflow showing the results of multiplexed immunofluorescence (mIF) imaging, cell segmentation, cell phenotyping, cellular neighborhood analysis, contour annotations, and multimodal data integration using SpatialData.

### Providing a versatile solution to support diverse spatial proteomics experiments

We built openDVP for compatibility with a wide range of imaging applications that support whole-slide images (WSI) and tissue microarrays (TMA) based on conventional widefield or confocal microscopy (H&E, IHC, IF, and mIF) (**Fig. 1b**). While WSI can provide a comprehensive view of large tissue areas, which are particularly important for heterogenous tissues, TMA applications support cohort-size spatial proteome analyses. Importantly, the imaging pipelines use BioFormats compatible file formats (e.g., OME-TIFF) and do not rely on any proprietary software or specific microscopy hardware for broad accessibility.

### Efficient and extensible image processing

Following image acquisition, the first step in DVP workflow is the processing of images, including illumination correction and stitching image tiles. OpenDVP builds on top of the established open-source image processing pipelines MCMICRO and SOPA (**Fig. 1a**). MCMICRO was built using Nextflow ^16^, following nf-core guidelines, to ensure high standards of modularity, consistency, and interoperability. SOPA is built on python-based Snakemake ^17^, enabling python-fluent users to add new modules or modify existing steps. We developed custom modules to expand segmentation masks by a defined number of pixels, quantify marker intensity quantiles per cell, and pyramidize images for smooth quality control. These open-source pipelines offer key advantages: (1) Broad compatibility, supporting most imaging formats and modalities. (2) Modular architecture, integrating latest benchmarked algorithms and machine learning models, such as Cellpose ^18^ and Mesmer ^19^. (3) Scalability, enabling efficient distribution of processes for stitching, registration and segmentation. For example, segmenting eleven WSI with a total of 6.34 million cells took less than two hours (**Extended Data Fig. 1a-c**). (4) Active maintenance and development by a growing community of bioimage analysts.

### Interactive quality control and marker thresholding for single-cell phenotyping

Visual quality control is critical in image processing pipelines to assess illumination correction, stitching and registration results, and segmentation accuracy. OpenDVP also provides demo datasets and Jupyter notebook-based tutorials to help users integrate and inspect results using two complementary interfaces, Napari and QuPath. We chose them for their ability to visualize larger-than-memory images and community activity in developing new features. To facilitate image-based cell phenotyping based on marker expression, we developed an interactive Napari plugin (**Fig. 1c**) (github.com/CosciaLab/napari-cell-gater). Our plugin enables users to visualize multichannel images, overlay segmentation masks, plot mutually exclusive markers (FACS analysis-like), and provide real-time feedback of cells being labelled positive or negative based on selected thresholds. Although more laborious, this approach gives users full autonomy in their filter strategy for the complex task of single-cell phenotyping. For transparency, our plugin generates summary tables with cutoff values for each marker-sample pair. Finally, our image-processing pipeline creates a filtered single-cell matrix used for supervised or unsupervised analysis to identify cell types, states, and cellular neighborhoods. These guide laser microdissection and exploratory LC-MS-based proteome analysis.

### A versatile interface between image analysis and laser microdissection

Laser microdissection is essential in the DVP workflow for isolating single cells or ROIs precisely and automatically. Segmentation-based cell/nucleus isolation provides the highest biological granularity, particularly relevant for single-cell applications or rare phenotypes. The isolation of larger ROIs (e.g., multi-cellular niches) represents a robust alternative to single-cell LMD cutting. To support both scenarios, we developed QuPath-to-LMD, an open-source web app for experimental design, contour export, and validation of LMD-ready masks, supporting the Leica LMD7 and MMI CellCut systems (**Fig. 1d**) (github.com/CosciaLab/Qupath_to_LMD). QuPath-to-LMD provides an interface between histopathology and omics-based profiling, as slide annotations can be directly transferred into LMD-ready contours for downstream (prote)omic analysis. We validated single-cell and ROI-based contour alignment on the LMD7 system using different magnifications (20x and 40x) and provided example images for selecting optimal tissue reference points (**Extended Data Fig. 1d-e**).

### Integrating imaging and MS-based proteomics data through the scverse ecosystem

DVP generates two types of proteomic data: imaging and LC-MS data. For data integration, visualization, and multi-modal analysis, openDVP’s python code uses scverse standards and the AnnData and SpatialData formats, ensuring compatibility with popular analysis packages such as scimap ^20^, scanpy ^21^, and spatialproteomics ^22^. To our knowledge, this framework represents the first instance of integrating mass spectrometry-based proteomic data into the scverse ecosystem, streamlining data analysis for large-scale spatial tissue proteomics. For example, through SpatialData adoption, users can overlay their images with cellular phenotype information (e.g., cell types and cellular neighborhoods) and matching quantitative proteome information. We developed a data analysis package to facilitate storing, integrating, visualizing, and analyzing spatial proteomics data, available for download from the PyPi registry (**Extended Data Fig. 1f**). A vignette of the openDVP workflow is shown in **Fig. 1e**.

### OpenDVP quantifies cell-type resolved proteomes of human placental tissue

Understanding biological foundations of human health and disease requires comprehensive characterization of individual cells, the fundamental units of life. Recent single-cell and spatial omics advancements have catalyzed large-scale initiatives, such as the Human Cell Atlas ^23^ and the LifeTime Initiative ^24^, which aim to systematically profile all human cell types across tissues to unravel relationships between cells, tissue organization, and function. However, certain cell types (e.g., cardiomyocytes, adipocytes, and neurons) pose analytical challenges due to their resistance to dissociation into single-cell suspensions. Their interconnectivity and/or fragility make them inaccessible for sorting-based single-cell profiling, and they are often morphologically too complex for accurate cell segmentation. One example is the syncytiotrophoblast (STB) of the human placenta, a syncytial layer derived from fused cytotrophoblasts (CTB) that lacks distinct cell boundaries (**Fig 2a**), making cell segmentation and single-cell-based laser microdissection challenging. Despite STB’s critical roles in nutrient exchange, hormone secretion, and immune modulation at the maternal-fetal interface ^25^, its syncytial nature has prohibited its comprehensive molecular profiling, particularly at the global proteomic level. With this in mind, we applied flashDVP as a simplified and image segmentation-free version of DVP. We performed four-color immunofluorescence (IF) staining of 5µm thick first trimester human placental FFPE sections mounted on PPS frame slides and performed centimeter-scale whole-slide imaging (**Fig 2b**).

**Fig. 2:**
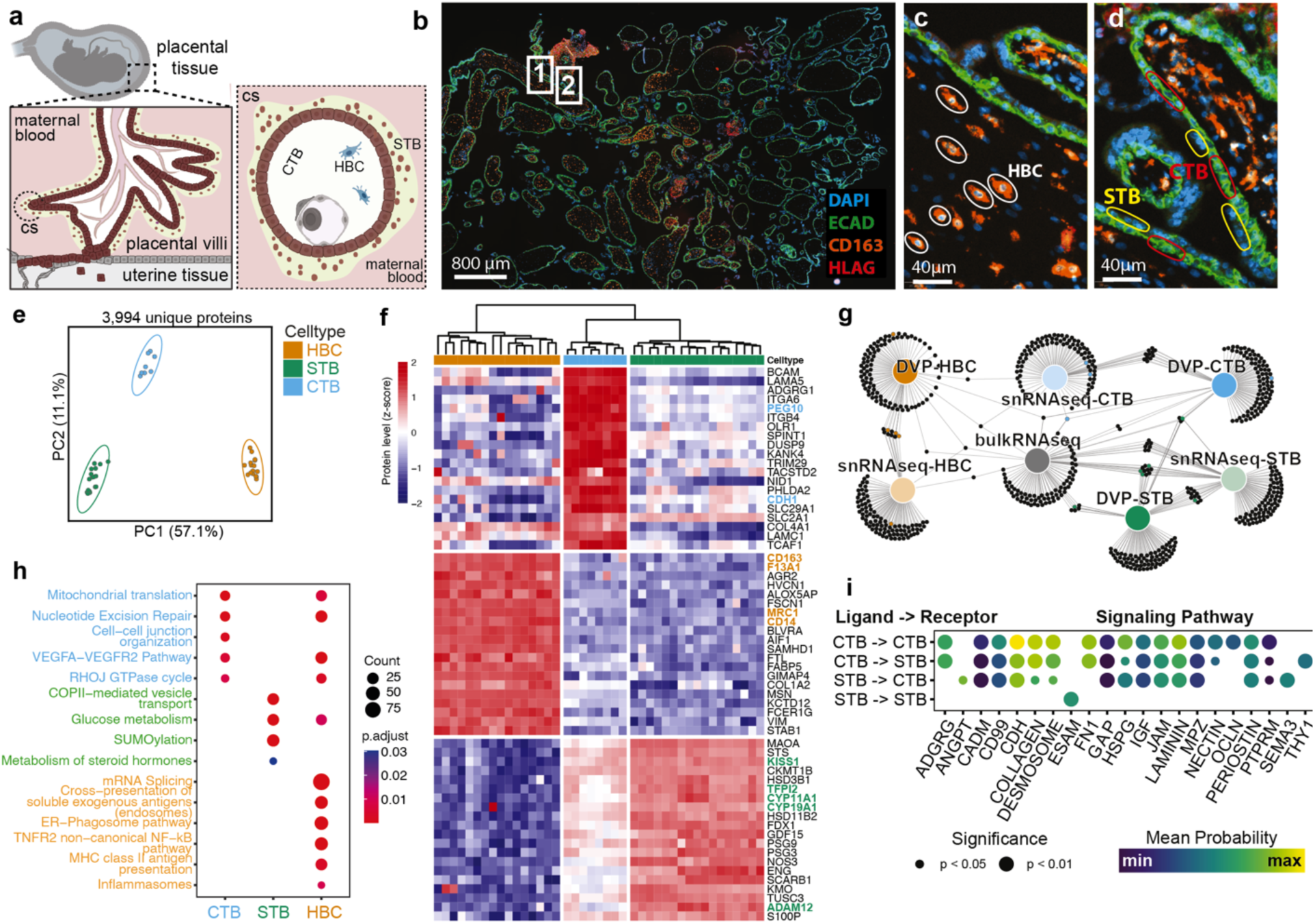
High-resolution cell-type resolved proteomics of human placental tissue. **(a)** Left: Schematic of placental villi samples and fetal structures at the maternal-fetal interface. Right: Cross-section showing cytotrophoblast (CTB) monolayer replenishing syncytiotrophoblast (STB) layer, with Hofbauer cells (HBC) in the stroma. cs, cross-section. **(b)** Representative immunofluorescence image of placental tissue stained for E-cadherin, CD163, HLA-G, and DNA (DAPI), used for cell type annotation, laser microdissection, and proteomic profiling. Box inserts correspond to panels c (1) and d (2). **(c,d)** Representative regions of interest manually selected in QuPath for isolation of immune (c) and trophoblast (d) compartments. CTB monolayer is defined by E-cadherin+ and HLAG-, whereas the multinucleated STB layer is E-cadherin- and HLAG- in the outer layer of the villi. CD163 expression marks macrophages (HBC). **(e)** Principal component analysis of cell type resolved proteomes from three placental tissues. Colors indicate cell types. Confidence ellipses (95% CI) computed from PC scores covariance. **(f)** Cell type-specific protein marker heatmap as inferred by ANOVA (FDR < 0.05) showing relative abundance profiles per replicate. Canonical markers for each phenotype highlighted in orange (HBC), green (STB), and light blue (CTB). Z-scored relative protein abundances are shown. Distance metric: “euclidean”, method: “complete”. **(g)** Comparison of flashDVP-generated phenotype-specific proteins and published genes marking CTB, STB, and HBC phenotypes. Shared and unique markers highlight the importance of proteomic data in complementing other omics. **(h)** Dotplot of enriched functional pathways of phenotype-separating clusters from (g). Enrichment based on ranked proteins using the Reactome database. Color shows adjusted p-values of overrepresented pathways and dot size indicates category sizes (“Count”). **(i)** Significant cell-cell communication in trophoblast compartment inferred using CellChat^36^. Dot colors show mean communication probability of ligand-receptor pairs in labelled pathway, size represents p-value from one-sided permutation test.

The placental barrier, appearing in tree-like villi structures, controls the nutrient and gas exchange between maternal and fetal blood. It consists of E-cadherin+ bipotential CTBs that fuse to form a multinucleated E-cadherin−, HLAG− STB layer that is in direct contact to maternal blood. HLAG+ trophoblasts constitute the invasive CTB derived lineage that anchor the placenta to the maternal uterine tissue (**Fig 2a**). CD163+ extraembryonic macrophages (Hofbauer cells, HBC) in the stroma play roles in tissue remodeling, immune surveillance, and support the formation and maintenance of fetal blood vessels in the villi ^26^. With QuPath, we manually annotated 150 cells per replicate for each cell type (STB, CTB, and HBC, **Fig 2c-d**) to reach an equal tissue amount per sample. LMD contours were created using the QuPath-to-LMD web app, transferred to a Leica LMD7 microscope for laser microdissection and processed using our ultralow-input tissue proteomics protocol ^27^. Samples were measured in label-free diaPASEF mode and analyzed with DIA-NN ^28^ in library-free mode. We identified 3,994 unique proteins across groups (**Extended Data Fig. 2a**), including canonical markers for each cell type (**Fig. 2d, Extended Data Fig. 2b-c**). Principal component analysis separated cell phenotypes regardless of tissue origin (**Fig. 2e**). Differential abundance analysis revealed cell type-specific markers that showed good concordance with public transcriptomics data ^29,30^ (**Fig. 2f-g, Extended Data Table 1**). Pathway analysis of protein clusters confirmed STB-specific functions in steroid hormone metabolism, vesicular transport, and energy demands (**Fig. 2h**). The regulation of SUMO proteins, linked to placental development and hypertension in pregnancy in bulk studies ^31,32^, was specific to STBs despite their lower abundance, underscoring the value of cell-type resolved proteomic analyses. CTB and HBC showed an enrichment in RhoJ GTPase and VEGF signaling pathways, while HBCs displayed immune regulation pathways, supporting their immunotolerant role. We found HBC-specific enrichment for proteins involved in mRNA splicing, aligning with the role of alternative splicing as an essential feature of placental dysfunction ^33^. Given the physical proximity between STBs and CTBs (**Fig. 2a, c**), cell communication analysis between these connected trophoblasts cells revealed high paracrine activity of both cell types, with autocrine communication strongest in the CTB monolayer (**Fig. 2i, Extended Data Table 1**). Strong signaling via cadherins (CDH), junctional adhesion molecules (JAM), fibronectins (FN1), and desmosomes was inferred, supporting the importance of cell adhesion integrity in fusion-competent CTBs. We identified robust communication involving collagens, laminins, and extracellular matrix proteins critical for implantation and placentation ^34^. Our data also revealed that endothelial cell-selective adhesion molecule (ESAM) as a unique and previously unreported autocrine signaling molecule of STBs. ESAM is an important component of tight junctions and a negative regulator of platelet activation; the latter is known to underpin pregnancy health and coexist with complications ^35^. These findings establish flashDVP as a powerful ‘fast-track’ approach for cell type and spatially resolved tissue proteomics.

### OpenDVP coupled to spatial transcriptomics uncovers niche-specific drug targets

We next tested whether flashDVP could facilitate cell-type resolved tissue proteomics by integrating additional spatial omics layers. We analyzed a serial FFPE tissue section from an early-stage lung adenocarcinoma that we previously characterized by spatial transcriptomics (ST, Nanostring CosMx) ^37^ (**Fig. 3a**). Using 960 cancer-related genes, we profiled over 340,000 cells, identifying 18 cell types and their spatial distribution. These rich data revealed diverse immune and tumor cell niches of therapeutic value, including a small tumor niche enriched for pro-metastatic mesenchymal tumor cells with high *NDRG1* and *LGALS1* expression, located near myofibroblasts and SPP1+ tumor-associated macrophages (TAMs) (**Fig. 3b**). Pseudotime analysis identified this ‘epithelial to mesenchymal (EMT)-niche’ as the potential start site of tumor invasion. We used the CosMx data to guide the collection of the EMT-niche tumor cells, and revealed their proteomes to understand their functional state and identify potential drug targets. IF staining of a serial tissue section against epithelial and mesenchymal markers, overlaid with CosMx phenotypes, confirmed the mesenchymal nature of EMT niche tumor cells on protein level (panCK+ and VIM+) (**Fig. 3c-d**). We next annotated multiple ROIs to proteomically profile tumor and stromal niches (**Extended Data Fig. 3a**). ROIs contained 50-100 cells (40,000 – 50,000 µm^2^, 5 µm thick). We quantified 4,138 and 4,976 proteins in stromal and tumor compartments respectively, totaling 5,956 proteins (**Fig. 3e, Extended Data Fig. 3b, Extended Data Table 2**).

**Fig. 3:**
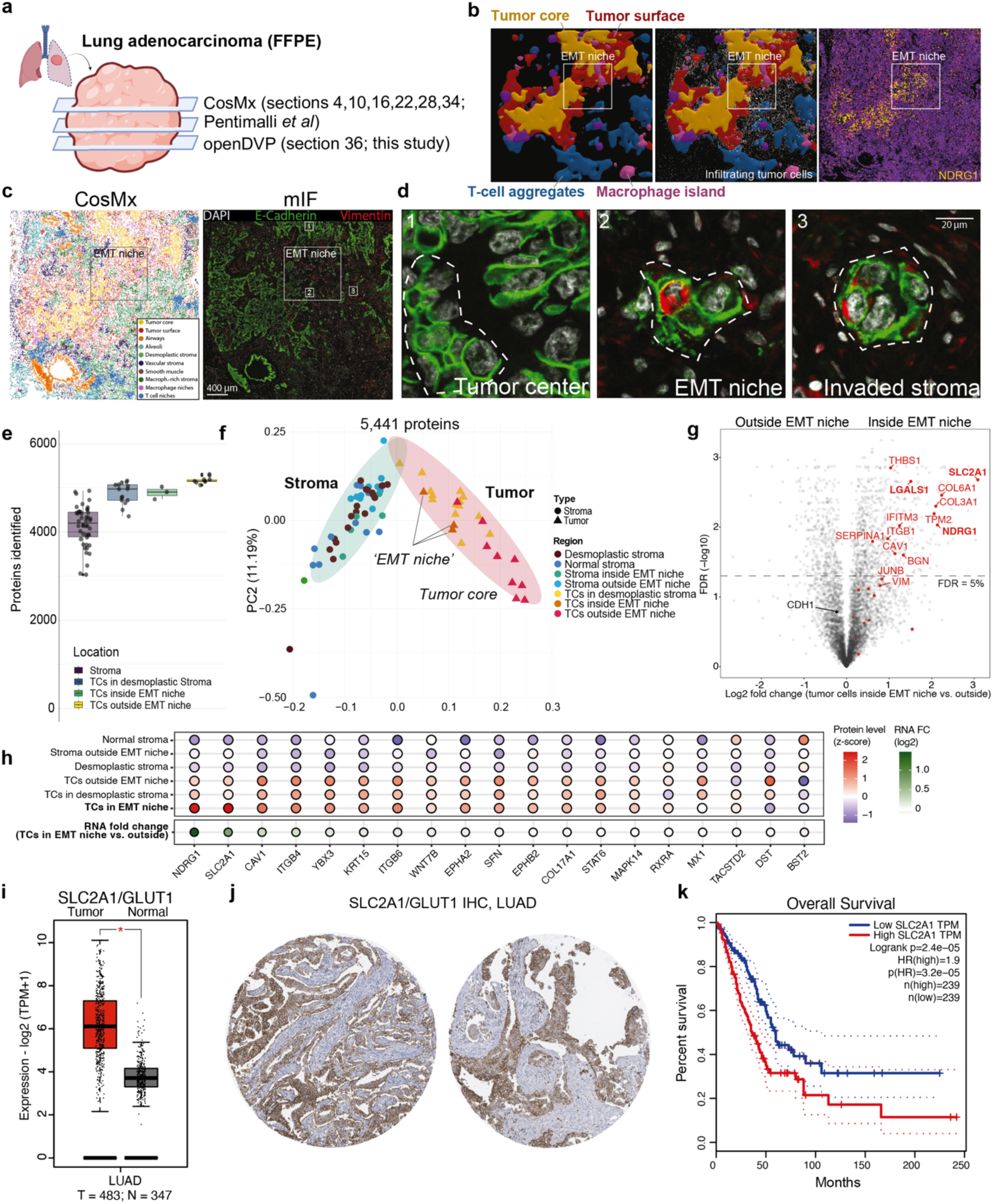
Cell-type resolved deep proteomic profiling of transcriptionally defined cellular niches. **(a)** Overview of 5 µm FFPE lung adenocarcinoma sections showing regions for CosMx spatial transcriptomics and DVP analysis. **(b)** 3D reconstruction of transcriptomically defined niches, showing tumor-infiltrating cells. NDRG1 RNA level was upregulated in tumor cells from the surface and EMT niche, as previously published by us ^37^. **(c)** Left: 2D multicellular niche map from CosMx data (Section 34^37^). Right: immunofluorescence of epithelial (E-cadherin, green) and mesenchymal (vimentin, red) markers in section 36. Insets 1–3 correspond to panel (d). Nuclei with DAPI (white). Scale bar: 400 µm. **(d)** Magnified views: [1] tumor center (E-cadherin+/Vimentin-), [2] EMT niche (E-cadherin+/Vimentin+, arrow), and [3] invading tumor cells (E-Cadherin+/Vimentin+, arrow). Scale bar: 20 µm. **(e)** Proteomics identified a mean of 4,138 proteins in stroma and 4,976 in tumor regions (50–100 cells/sample). **(f)** Principal component analysis of 5,441 proteins shows a separation of tumor and stroma samples along PC1. EMT niche tumor samples cluster near invasive and stromal samples. **(g)** Volcano plot showing differential protein abundance in tumor cells inside vs. outside the EMT niche. CosMx-upregulated genes ^37^ are shown in red. Proteins above the dashed line have a permutation-based false-discovery rate (FDR) below 0.05. **(h)** Relative protein levels of top upregulated genes in EMT niche tumor cells. **(i)** Mean RNA expression of *SLC2A1*/*GLUT1*, the most EMT-niche upregulated tumor protein in the proteomics data, in lung cancer cohort (n = 483 tumor, 347 healthy; TCGA LUAD) shows higher expression in cancer tissues ^41^. **(j)** GLUT1 IHC staining of lung adenocarcinoma tissues confirmed tumor-specific expression. Images (left: ID 426; right: ID 1303) from human protein atlas ^43^. **(k)** Kaplan–Meier analysis showed lower overall survival in patients with high *SLC2A1*/*GLUT1* RNA expression, grouped by median. For boxplots in **(e)** and **(i)** the central lines indicate the median, the box the interquartile range, whiskers extend to (1.5*IQR) of the box.

Group replicates clustered together and showed high proteome correlations (Pearson’s r > 0.9, **Extended Data Fig. 3c**). PCA analysis showed EMT-niche tumor cells and stroma-infiltrating tumor cells were closer to stromal samples in component 1, reflecting their mesenchymal phenotype (**Fig. 3f**). Integration with CosMx data confirmed that tumor cell-intrinsic signatures dominated the EMT-niche tumor proteomes (**Fig. 3g-h**). Proteomics further validated the CosMx and IF data showing panCK+, VIM+, and E-cadherin-tumor cells in the EMT-niche and upregulated NDRG1 and LGALS1 protein levels. Many genes identified by ST were also significantly upregulated at the protein level in EMT niche tumor cells, emphasizing the value of spatial niche information in 3D for aligning and integrating tissue sections for multi-omics analyses (**Fig. 3a**). However, MS-based proteomics captured higher dynamic ranges of analyte abundances, resulting in different gene/protein rankings and drug target prioritizations (**Fig. 3g-h, Extended Data Fig. 3d**). We identified 180 FDA-approved drug targets of which 16 cancer-related genes were significantly upregulated in EMT niche tumor cells (**Extended Data Fig. 3e, Extended Data Table 2**). For example, *PDGFRB*, an actionable gene for imatinib, a receptor tyrosine kinase inhibitor approved for various malignancies including gastrointestinal stromal tumors and myeloproliferative neoplasms ^38^. SLC2A1/GLUT1 was the most up-regulated protein in EMT niche tumor cells. Consistently, pathway analysis revealed high glycolysis expression in EMT niche tumor cells, along with hypoxia, angiogenesis, and EMT pathways (**Extended Data Fig. 3f**). As targeting glucose metabolism is currently under clinical investigation for cancer therapy ^39,40^, we assessed the tumor-specific expression of GLUT1 and its association with patient outcomes. We validated high tumoral GLUT1 expression in a large-scale transcriptome study of 483 lung adenocarcinomas and 347 control samples ^41,42^ (**Fig. 3i)** as well as on IHC data obtained from the human protein atlas ^43^ (**Fig. 3j)**. High *GLUT1* expression was a strong prognostic factor associated with poor patient outcome, as revealed by the significantly different overall survival (*p* = 0.0003, n=239 per group) (**Fig. 3k**). Notably, re-evaluation of the spatial transcriptomic data not only confirmed the tumor-specific expression of *GLUT1*, but also emphasized its strong EMT-niche specificity (**Extended Data Fig. 3g**). Together, these results underline the benefit of cellular niche-guided spatial multiomics as a promising approach for identifying and validating niche-specific, personalized drug targets.

### OpenDVP reveals tumor microenvironment remodeling in primary and relapsed TNBC

Current DVP implementations enable the linkage of visually and machine learning defined tissue phenotypes to quantitative proteomic profiles but involve trade-offs between spatial resolution, proteome depth, and experimental throughput. Our original DVP concept ^8^ achieves workflow robustness and high proteome coverage by pooling phenotypically coherent cell populations, whereas single-cell DVP (scDVP) ^44^ maximizes spatial resolution at the expense of analysis depth and ease of use. These limitations motivate alternative strategies that preserve spatial interpretability while substantially increasing throughput and discovery potential.

Recent advances in spatial biology have established cellular neighborhoods (CNs) as fundamental organizational units that shape tissue function, disease progression, and therapeutic response ^45–47^. Triple-negative breast cancer (TNBC), the most aggressive breast cancer subtype ^48^, exemplifies how CN organization within the tumor microenvironment (TME) correlates with clinical outcomes ^49^, including critical roles for macrophage- and fibroblast-enriched niches ^50–52^. Resolving CN-specific proteomes therefore offers an opportunity to identify molecular mediators of cell–cell interactions and disease progression. To enable CN-resolved proteome profiling, we first established an optimized protocol for centimeter-scale whole-slide multiplex immunofluorescence (mIF) imaging on PPS membrane slides, using a nine-marker panel for this proof-of-concept study (**Fig. 4a-b**). We analyzed paired TNBC tissue sections from the primary surgical resection and a matched local relapse obtained two years after adjuvant chemotherapy. The primary specimen contained extensive ductal carcinoma in situ (DCIS; 38 mm) with focal invasion into the surrounding stroma, whereas the relapse consisted of a large (45 mm) invasive carcinoma without residual DCIS (**Fig. 4c**). Each whole-slide image stack (>150 GB) was processed using MCMICRO and SOPA for cell segmentation, phenotyping, and neighborhood analysis, with matched H&E sections providing additional morphological context (**Extended Data Fig. 1a-c**). We segmented 610,182 and 1,005,051 cells from the primary and relapse samples, respectively, identifying five major cell types —tumor, T cells, B cells, macrophages, and stromal cells (**Fig. 4c–e; Extended Data Fig. 4a-d**). While panCK⁺ tumor cells dominated both tissues, marked shifts in immune and stromal composition were observed (**Fig. 4f; Extended Data Table 3**). Compared with the primary tumor, the relapse exhibited pronounced depletion of B and CD8⁺ T cells alongside a 16-fold increase in CD68⁺ macrophages and a fourfold expansion of VIM⁺ stromal cells, consistent with immune evasion and tumor progression ^53,54^. Cellular neighborhood analysis using scimap’s spatialLDA identified seven recurrent CNs (CN0–6) with distinct cellular compositions (**Fig. 4g; Extended Data Table 3**). Several CNs showed strong abundance differences between the two samples (**Fig. 4g–i**), accompanied by spatial reorganization of immune cells. Proximity analysis revealed that macrophages were located closer to tumor cells in the relapse, whereas lymphocytes preferentially surrounded DCIS regions in the primary tumor (**Fig. 4j**), indicating a transition from a lymphocyte-rich peritumoral niche to a macrophage-dominated intratumoral microenvironment.

**Fig. 4.**
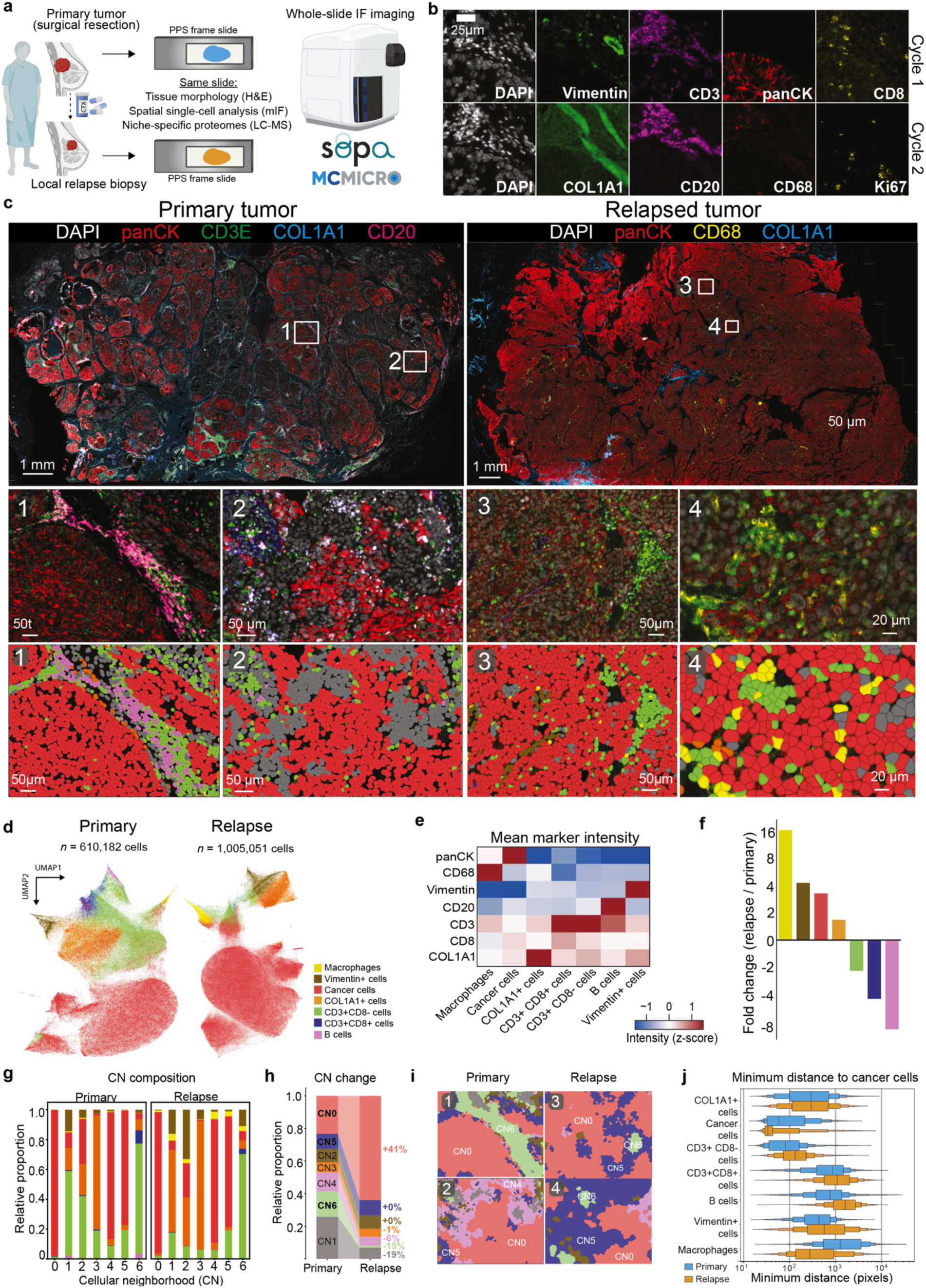
Image acquisition and analysis of primary and recurrent triple-negative breast cancer (TNBC). **(a)** FFPE 5 µm sections from primary tumor and same-patient post-chemotherapy relapse sample were mounted on PPS frame slides. **(b)** Slides underwent multiplex immunofluorescence (mIF) staining and processing using MCMICRO pipeline with SOPA. **(c)** Whole-slide mIF images of samples with zoomed regions show overlaid cell phenotype information. **(d)** UMAP embeddings of segmented cells from each slide, based on morphology and marker intensity (excluding nuclear stains and background); unidentified cells excluded; n_neighbors = 25. **(e)** Heatmap of z-scored mean marker intensities across phenotyped cells. Phenotyping performed by thresholding markers per slide, rescaling intensities, and scimap phenotype function. **(f)** Fold change in cell type counts between recurrent and primary samples. **(g)** Relative composition of cellular neighborhoods per sample excluding unidentified cells. h) Selected regions with cellular neighborhoods overlaid. **(i)** Stacked bar plots showing cellular neighborhood frequency in each sample, calculated using scimap’s spatialLDA function. **(j)** Boxen plots of minimum distance from cells to nearest cancer cell. The central line indicates the median, while the first box represents the interquartile range (IQR, 25th to 75th percentiles), each subsequent level of boxes corresponds to half the previous number of datapoints.

### Cellular neighborhood–guided DVP enhances proteome coverage and scalability

Motivated by these spatial differences, we hypothesized that cellular neighborhood–guided DVP (cnDVP) could enable scalable spatial proteomics while preserving the core DVP principle of linking visual phenotypes to quantitative proteomic profiles. To systematically evaluate this approach, we compared cnDVP with conventional DVP ^8^ and scDVP ^44^ using matched regions from the primary and relapse samples. To minimize spatial confounders, we profiled directly adjacent cells of identical phenotype and CN assignment (**Fig. 4e,g**; **Fig. 5a**). For DVP, we excised 100 individual cell shapes per well, whereas cnDVP required only a single shape encompassing 100 phenotyped cells, resulting in an approximately 100-fold faster tissue collection. For scDVP, we collected individual tumor or immune cells. As cell size largely determines proteome coverage ^55^, scDVP has remained challenging for small cell types such as immune cells (**Extended Data Fig. 5a**). From 50 single immune cells (CD3+/CD8-lymphocytes or CD68+ macrophages) only three samples remained after filtering for more than 400 proteins. For cancer cells, we identified up to 1,500 proteins per shape (**Fig. 5b**). CnDVP achieved the highest proteome coverage in both tumor and immune niches, with a 26% increase in protein identifications relative to DVP and the largest number of uniquely detected proteins (**Fig. 5b-c**). The majority of proteins were consistently detected across all approaches (**Extended Data Fig. 5b**), and tissue- and niche-specific quantitative signatures were highly correlated (**Fig. 5d-e, Extended Data Fig. 5c-e**). These results establish cnDVP as an efficient and generalizable strategy that substantially improves throughput while largely preserving cell-type specificity and spatial resolution.

**Figure 5:**
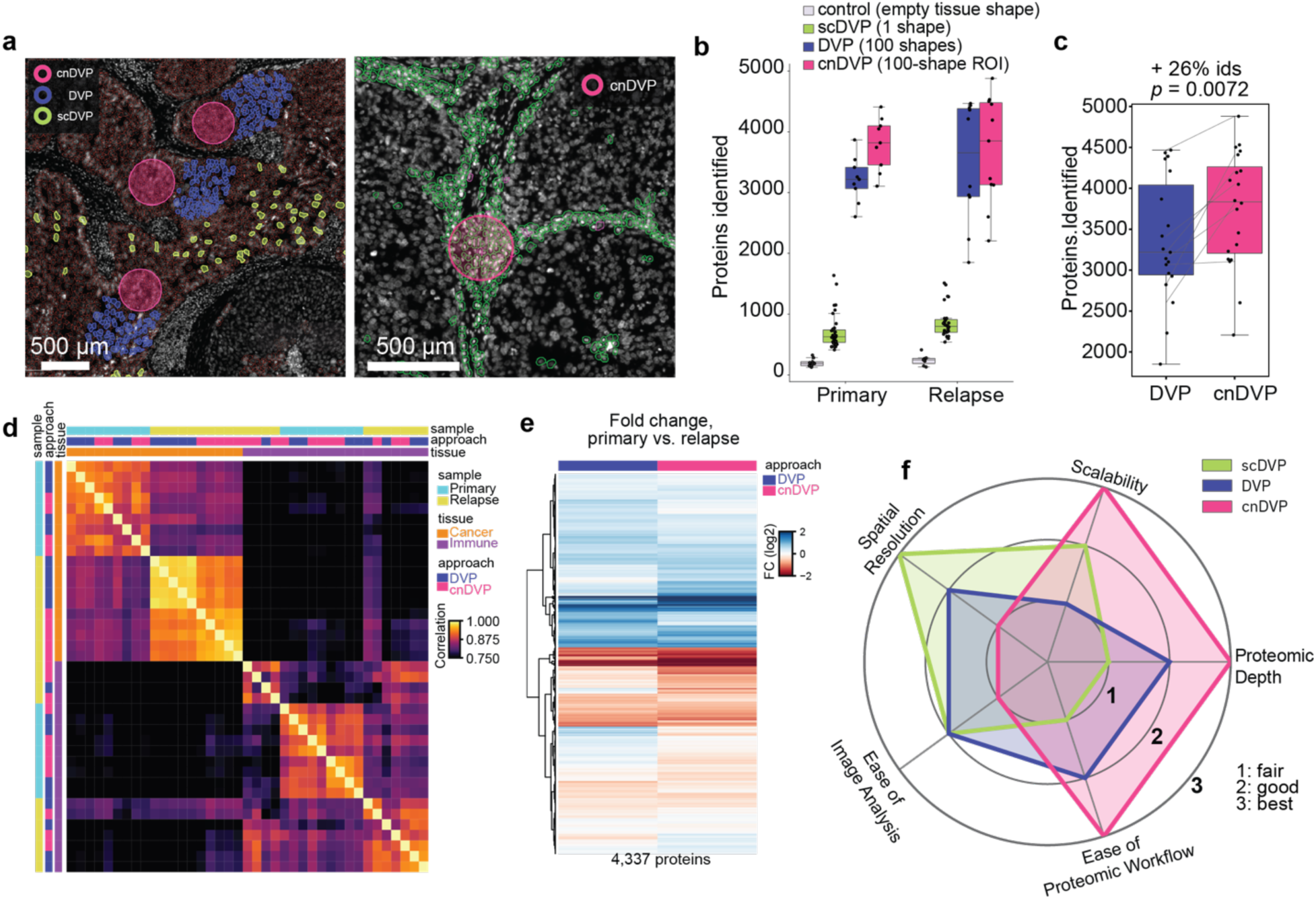
Cellular neighborhood guided DVP improves proteomic depth and throughput. **(a)** Images of primary TNBC tissue, showing the nuclear stain with segmentation masks for phenotyped cancer cells with red outlines on the left and CD3+ cells with green outlines on the right. Overlayed are collection annotations for the three DVP approaches. In magenta cnDVP, in blue DVP, and green for scDVP. **(b)** Boxplots with number proteins identified with DIANN search across DVP approaches. **(c)** Boxplot with number of proteins identified in matching cancer-specific samples analyzed by DVP and cnDVP. Lines denote spatial co-localization, showing an increase of proteome coverage using the cnDVP approach. P-value is based on a two-sided paired T-test between paired samples. For both b and c, the central lines indicate the median, while the box represents the interquartile range (IQR, 25th to 75th percentiles), whiskers extend to 1.5* IQR of the box **(d)** Proteome correlation heatmap using Spearman correlations for all immune and cancer samples analyzed with DVP and cnDVP. Metric: “euclidean” and method: “ward” **(e)** Heatmap of log2FC values for cancer samples between the primary and relapse breast cancer tissues across DVP and cnDVP approaches. Note, similar fold changes were retrieved for both approaches. **(f)** The radar plot summarizes the key differences between three DVP approaches; scDVP (light green), DVP (blue), and cnDVP (pink); across five key properties: Scalability, Proteomic Depth, Ease of Proteomic Workflow, Ease of Image Analysis, and Spatial Resolution. The radial axes represent a qualitative ranking scale where 1 = “fair”, 2 = “good”, and 3 = “best”.

Conceptually, cnDVP expands the DVP method family implemented in openDVP, enabling users to select among complementary profiling strategies based on tissue architecture, biological question, and experimental constraints (**Fig. 5f**).

### CN-guided DVP delineates spatial tumor heterogeneity and molecular drivers of relapse

To investigate how CN-resolved proteomes reflect tumor progression and treatment resistance, we performed CN-guided LMD sampling for exploratory proteomics (**Fig. 6a**), focusing on tumor-specific (CN0), tumor–immune interface (CN5), and immune-enriched (CN6) regions. Regions of interest were distributed across tissue sections to capture spatial heterogeneity (**Fig. 6b**), yielding 203 low-input samples after QC filtering, and processed within one week using an advanced throughput (80 samples per day) DIA-based workflow. We quantified ∼4,800 proteins in tumor-specific CNs and 3,500–4,000 proteins in immune-enriched niches (**Extended Data Fig. 6a-b**). Proteomes exhibited strong CN specificity, with shared core proteins and niche-restricted signatures (**Fig. 6c**). Integration with SpatialData and public scRNA-seq datasets ^56^ confirmed epithelial tumor signatures in CN0 and immune programs in CN6 (**Fig. 6d**). Principal component analysis separated proteomes by tissue origin and spatial niche (**Fig. 6e-f**), and unsupervised clustering identified distinct proteomic programs associated with primary versus relapsed tumor samples (**Fig. 6g-h**). Tumor-specific CNs showed significantly higher proteomic heterogeneity in the primary sample than in the relapse, as reflected by lower global correlation, increased protein variability, and stronger spatial autocorrelation (**Fig. 6i,j; Extended Data Fig. 6c–e**). This observation also transcended to single tumor shapes (**Extended Data Fig. 6e**), possibly reflecting reduced intratumoral diversity through clonal selection following chemotherapy ^57,58^.

**Figure 6.**
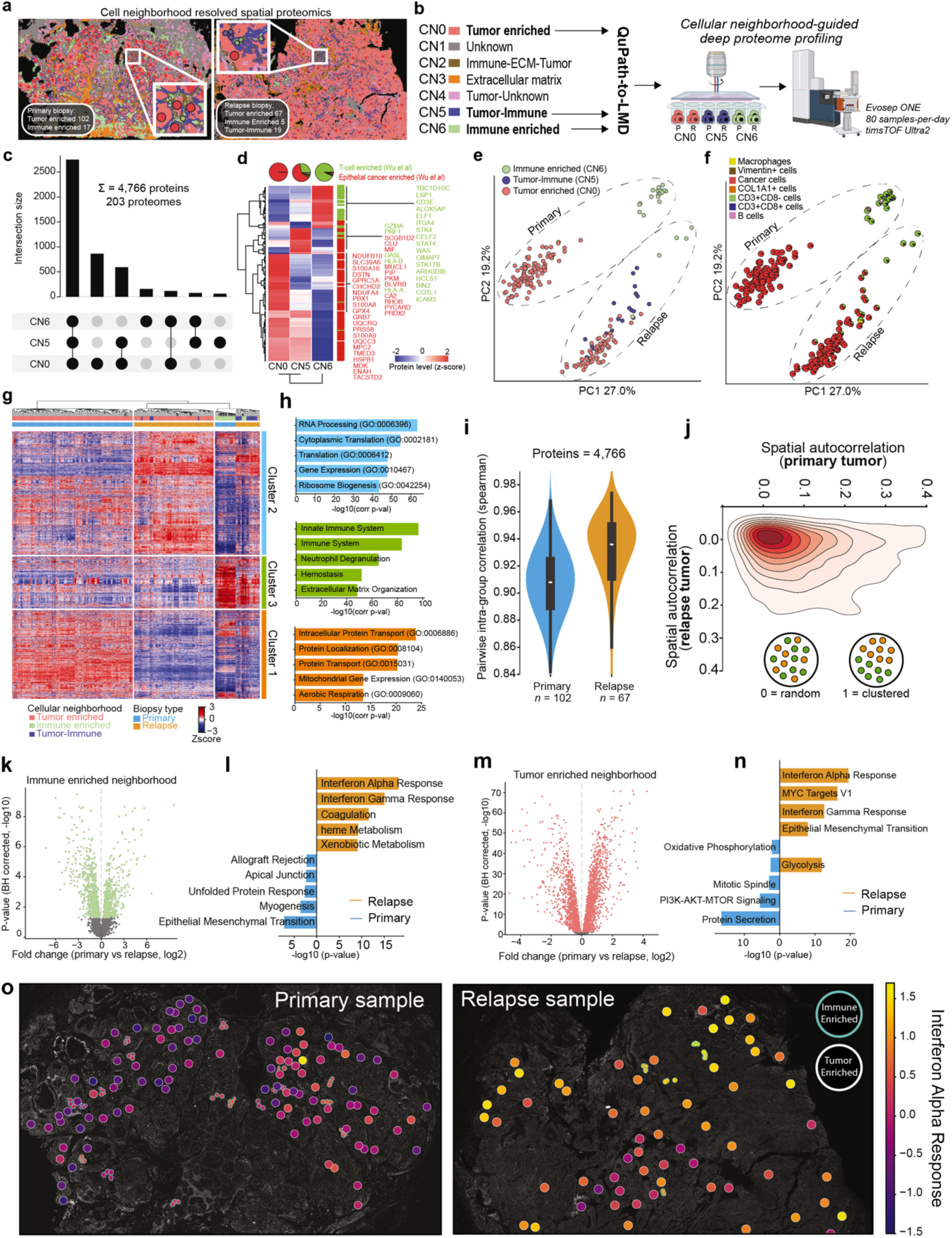
Spatial proteomic analysis of triple-negative breast cancer (TNBC) samples comparing primary and recurrent disease. **(a)** Whole-slide overlay of cellular neighborhood (CN) annotations with contours for laser microdissection-guided proteomics. Sample collection summaries are shown in boxes. **(b)** Three selected CNs for proteomics: Tumor Enriched, Immune Enriched, and Tumor-Immune. Regions were collected using laser microdissection and analyzed using a rapid (80 samples-per-day) LC-MS strategy. **(c)** UpSet plot showing protein identification overlap across CNs. **(d)** Heatmap showing signature markers from a recent RNAseq study ^56^. Protein levels were z-scored, averaged per CN, and subjected to unsupervised hierarchical clustering (method: “ward”, distance metric: “euclidean”). **(e)** Principal component analysis (PCA) of all samples, colored by CN. **(f)** PCA embedding with samples as pie charts reflecting cell phenotype proportions derived from image analysis. Note, both PCA plots show clustering by sample type (primary vs. relapse), followed by CN. **(g)** Heatmap displaying 4,570 proteins across 203 samples, clustered hierarchically (method: “ward”, distance metric: “euclidean”). Proteins filtered for 70% valid values in any CN group. **(h)** Protein clusters analyzed using EnrichR pathway enrichment using all identified proteins as background. **(i)** Violin plots showing pairwise intragroup correlations, highlighting greater proteomic variability in the primary tumor. P-value <0.001, Mann-Whitney two-sided test. Central lines indicate the median, box represents the interquartile range, whiskers extend to 1.5* IQR of the box **(j)** Density plot of Moran’s I spatial autocorrelation for proteins across samples, showing greater spatial variability in the primary tissue. P-value <0.001, Mann-Whitney two-sided test. **(k)** Volcano plot showing regulated proteins between immune-enriched CNs of primary and relapse tumors, FDR cut-off of 5%. **(l)** Barplot showing enriched MsigDB hallmarks between primary and relapse immune-enriched samples. **(m)** Volcano plot showing regulated proteins between tumor-enriched CNs of primary and relapse samples, FDR cut-off = 5%. **(n)** Barplot showing enriched MsigDB hallmarks between primary and relapse tumor enriched samples. **(o)** Overlay of interferon alpha response pathway (Hallmarks) showing z-scored pathway levels of tumor and immune-enriched samples. Note, the relapse sample featured consistently higher IFN levels across tissue. Large circles represent tumor specific CNs (white outline) and small circles immune specific CNs (cyan outline).

Differential abundance analysis revealed pronounced niche-specific alterations in the relapse sample. Immune niches exhibited upregulation of interferon-γ signaling, complement, and heme metabolism pathways (**Fig. 6k-l**), alongside enrichment of myeloid/macrophage programs (**Extended Data Fig. 6f**). Tumor niches displayed a metabolic shift toward glycolysis and activation of EMT, MYC, and interferon signaling pathways (**Fig. 6n**), hallmark features of chemoresistance and progressive disease^59,60^. Spatial mapping revealed widespread interferon-high tumor phenotypes in the relapse but not in the primary tumor (**Fig. 6o**), consistent with molecular features of inflammatory TNBC and tumor-immune evasion ^61,62^.

Together, these results demonstrate how openDVP integrates whole-slide imaging, cellular neighborhood analysis, and exploratory proteomics to resolve spatial tumor heterogeneity and uncover potential drivers of disease progression.

## Discussion

OpenDVP is the first Python-based foundational platform designed to make deep visual proteomics broadly accessible and community driven. Built on the scverse ecosystem, it adheres to FAIR principles by ensuring that processed data are findable, accessible, interoperable, and reusable. Its modular architecture enables seamless integration of established tools for image segmentation, cell phenotyping, and spatial analysis, including MCMICRO, QuPath, and Napari, thereby supporting flexible and reproducible end-to-end DVP workflows.

For broad accessibility, openDVP provides two complementary workflow options. The first, flashDVP, implements a simplified ‘annotate–collect–measure’ strategy that enables rapid spatial discovery within days. This approach is particularly well suited for exploratory studies of acellular regions such as the extracellular matrix, or tissue contexts where single-cell phenotyping is less applicable. Applied to human placental tissue, flashDVP enabled comprehensive cell-type–resolved proteomic profiling, identifying pathways critical for placental biology. In an aggressive lung cancer previously analyzed by spatial transcriptomics, flashDVP confirmed transcriptional findings at the protein level and uncovered additional niche-specific drug target candidates, illustrating the complementarity of multimodal spatial omics.

The second workflow represents an AI-powered DVP pipeline optimized for high-performance computing and large-scale imaging data. This pipeline integrates state-of-the-art tools for memory-efficient single-cell and spatial analyses and was applied to study tumor relapse in triple-negative breast cancer. While our original DVP workflow and single-cell DVP offer maximal biological granularity, indispensable for certain biomedical applications ^44,63,64^ , they remain technically challenging due to gravity-based collection constraints, contour alignment errors, and potential cell-type admixture during single-cell LMD. These limitations motivated the development of an alternative strategy that balances biological resolution with scalability, robustness, and experimental efficiency. By operating at the level of cellular neighborhoods, cnDVP overcomes throughput limitations of existing DVP workflows while preserving spatial and cell-type specificity. We anticipate cnDVP as particularly useful for homotypic niches. For heterotypic niches, such as the case for tumor and immune cell co-localization, cnDVP could offer a direct and quantitative approach for protein-centric ligand-receptor analyses. Applied to a TNBC relapse case, cnDVP revealed extensive tumor microenvironment remodeling and pronounced proteomic differences between primary and relapse tumors. Our data revealed elevated interferon signaling, EMT, and glycolytic programs in relapse-associated niches - features linked to tumor progression and chemoresistance and potentially targetable through JAK/STAT inhibition or metabolic interventions ^40,65,66^.

Looking ahead, we anticipate that cellular neighborhood–guided proteome profiling will be particularly powerful when combined with higher-plex DVP assays ^67^ (e.g., 20–30 markers), substantially reducing undetected phenotypes and enabling finer-resolution dissection of tissue architecture and disease-associated states.

In summary, openDVP represents a significant methodological advance in spatial proteomics by providing an open, extensible, and scalable platform for deep visual proteomics. As imaging, AI, and proteomics technologies continue to mature, widespread adoption of openDVP across tissues and disease contexts has the potential to accelerate biological discovery, enable systematic biomarker identification, and uncover novel therapeutic targets. Community-driven efforts built on openDVP will further facilitate the development of centralized spatial proteomics resources that integrate antibody- and mass spectrometry–based data, ultimately advancing our understanding of tissue biology in health and disease.

## Acknowledgements

We thank our colleagues at the Max Delbrück Center (MDC) and Charité for their support and fruitful discussions. Furthermore, we acknowledge the MDC technology platform ‘Proteomics’ and ‘Advanced light microscopy’ for their great support. Gina Dörpholz, Pia Larsen and Jeannine Engel for administrative support and Janett König for laboratory support. Wouter-Michiel Vierdag for help with Napari plugin development and SpatialData. The authors also gratefully appreciate Martin Gauster (Medical University of Graz) and Andreas Glasner (Femina-Med) for patient recruitment collection of first-trimester human placental tissue (Medical University of Graz). J.N. and F.C. and S.S. acknowledge funding support by the Federal Ministry of Education and Research (BMBF), as part of the National Research Initiatives for Mass Spectrometry in Systems Medicine, under grant agreement No. 161L0222 and 03LW0239K. F.H. and F.C. were supported by Deutsche Forschungsgemeinschaft (HE 6249/5-3). This project received funding from the European Research Council (ERC) under the European Union’s Horizon 2020 research and innovation program (grant agreement No. 101115681) and support by the ERC (ERC starting grant). S.Fl. was supported by grant 01ZX1917A from the Federal Ministry for Education and Research (BMBF) through the e:Med Initiative for Systems Medicine.

## Author contributions

Conceptualization: J.N. and F.C.

Methodology: J.N., S.F., M.K., and M.T.

Resources: T.P., S.Fl., S.S., F.H., and F.K.

Experiments, J.N., S.F., M.K., and D.S.V.

Data curation: J.N., S.F., M.T., and D.S.V.

Data analysis: J.N., S.F., D.S.V., T.P., and F.C.

Figures: J.N., S.F., D.S.V., T.P., and F.C.

Project administration: J.N. and F.C. Supervision: F.H, S.Fl., N.R., and F.C.

Funding acquisition: N.R., F.H., F.K, and F.C.

Writing of the original draft: F.C.

All authors have reviewed and edited the manuscript.

## Methods

We structured the Methods section by analytical process to clarify workflow differences among the three exemplary applications. Detailed information on data sources, image preprocessing, proteomic analysis, and all other relevant settings is provided under dedicated subtitles for each vignette within these process-based descriptions.

### Sample collection and patient cohort

#### Placenta samples

Placental tissue was collected from electively terminated pregnancies (gestational age, 7 – 11 weeks) with informed consent. Exclusion criteria were maternal age under 18 years, BMI >25 kg/m2, and maternal pathologies. Ethical approval was obtained from Medical University Graz Ethics Committee (31-019 ex18/19). After surgical extraction, tissue was stored at 4°C in culture medium (DMEM/F12 1:1, 1 g/dL glucose) and processed within 4 h. Villous tissue was rinsed in cold (4°C) 0.9% NaCl solution to remove blood, fixed with 10% formalin, paraffin embedded (FFPE), and dehydrated per standard protocols.

#### Lung cancer sample

We analyzed the primary tumor of a non-small cell lung cancer (NSCLC) patient. The patient, a 63-year-old woman with a 40 pack-year smoking history and no physical limitations (ECOG performance status 0), was diagnosed in March 2020 with a metabolically active tumor at the apex of the right upper lobe on positron emission tomography (PET) imaging. A transbronchial lung biopsy confirmed a TTF1-positive lung adenocarcinoma (LUAD) with acinar growth pattern. Two months after diagnosis, the patient underwent right upper lobectomy. Gross examination was performed according to standardized protocols. The specimen was fixed in 10% buffered formalin and paraffin-embedded (FFPE). Histological examination, including diagnosis, tumor grading, pTNM classification, angioinvasion, lymphatic invasion, and tumor stage, was performed according to the 8th edition of the TNM classification (AJCC). Following diagnosis, the FFPE tissue block was stored at room temperature in the archive of the Institute of Pathology at the Charité University Hospital, Campus Mitte. The study was performed according to the ethical principles for medical research of the Declaration of Helsinki, and approval was approved by the Ethics Committee of the Charité University Medical Department in Berlin (EA4/243/21).

A comprehensive 3D spatial transcriptomic analysis of this tumor has been described recently based on 34 5-μm-thick consecutive sections (sections 4, 10, 16, 22, 28, and 34) ^37^ and a section-to-section distance of 30 μm. From the same tumor block, we profiled section 36 using proteomics after image registration in QuPath.

#### Triple negative breast cancer samples

Archival leftover diagnostic tissue from the tumor resection specimen and a specimen obtained two years later after relapse was selected from the archives of the Institute of Pathology, Charité Universitätsmedizin Berlin. At the time of diagnosis and primary resection, the patient had carcinoma in situ (DCIS), 38 mm in diameter, with multifocal stromal invasion. The multifocal invasive tumor had a total diameter of 9 mm, lacked expression of estrogen and progesterone receptors, and had an Her2-score of 1+, considered equivalent to a lack of HER2-gene amplification. The diagnosis was multifocal invasive triple-negative breast cancer (stage pT1b (m)). In 2022, the patient experienced a relapse with a diameter of 34 mm and ulceration of the skin (stage yrpT4b). The patient provided written consent for the use of leftover diagnostic material for research purposes. The project was approved by the IRB of Charité (“Ethics Committee”), project number EA1/253/19.

#### Membrane slide preparation

Prior to tissue mounting, slides were washed with 70% EtOH for 5min, rinsed in ddH2O and incubated with poly-L-lysine at 0.01% (w/v) (Sigma-Aldrich, cat.no. P1524) for 10min to optimize tissue adherence. Slides were then dried overnight at room temperature. All samples were sectioned at a thickness of 5 µm and mounted on metal frame PPS membrane slides (Leica, cat.no. 11600294), followed by overnight drying at 37 °C. Immediately prior to deparaffinization, slides were incubated at 60 °C for 30 min.

### Cyclic and non-cyclic immunofluorescence staining and imaging

#### Placenta tissue

Placenta tissue sections were deparaffinized, and subjected to antigen retrieval in Tris/EDTA (pH 9) at 93 °C for 20 min. After cooling (20 min), slides were placed in warm distilled water (5 min) and cooled again (5 min). Sections were washed with PBST (PBS + 0.1% Tween 20) and blocked with Ultra V Block (20 min, RT). For triple staining (CD163, CD31, E-cadherin), primary antibodies (Table 1) were diluted in PBST + 1% normal goat serum (NGS) and incubated overnight at 4°C. fter washing, secondary antibodies in PBST + 1% NGS (mouse IgG-AF647 and rabbit IgG-AF488, Table 1) were incubated for 1 h at RT in the dark. Slides were washed and mounted with SlowFade Diamond containing DAPI. Rabbit immunoglobulin and mouse IgG controls showed no stains. Imaging was performed using Zeiss Axioscan 7.

#### Lung and breast cancer tissue

Breast cancer and lung cancer tissue sections were deparaffinized and rehydrated (2x 5 min in Neo-Clear, 2x 2 min in 99% EtOH, 1x 2 min in 80% EtOH, 1x 2 min in 70% EtOH, 3x 1 min in 1x PBS). For decrosslinking, heat-mediated antigen retrieval was performed in pH 9 Tris buffer (Dako, #K800421-2, 1:50 in ddH_2_O) in a steamer for 30min. Sections were washed in 1x PBS and pre-quenched in 4.5% H_2_O_2_ (in 25mM NaOH in PBS) for 2x 30 min. Cover glasses were mounted with 10% glycerol (in PBS) and the first image acquisition for background subtraction was performed using the Zeiss Axioscan 7 Slidescanner with a 20x/0.5NA objective at 2 × 2 binning. Slides were soaked in PBS for cover glass detachment for 5 min and then washed 3x 5 min in PBS to remove residual glycerol. Tissues were then blocked for 30 min in 3% BSA (in PBS, Serva #11948.01) at room temperature, followed by antibody staining in a humid chamber at 4 °C overnight. Antibodies are listed in the table below, all diluted 1:50 in the blocking buffer. The next day, sections were washed in PBS, mounted, and imaged with similar settings, while exposure times were adjusted to the fluorescent signal. For cyclic (multiplex) immunofluorescence of breast cancer samples, cover glasses were removed as before, sections were washed, and fluorescent signal was bleached as previously described for 30 min. Sections were washed 3x 5 min in PBS, and the second cycle of antibody staining was performed.

Afterwards, sections were incubated for 3 min in hematoxylin, 10 min in tap water, dipped twice in ddH_2_O, 30sec in eosin, dipped twice in ddH_2_O, and dehydrated by dipping in increasing concentrations of EtOH (70-99%) for visible staining.

Immunofluorescent staining of lung cancer samples was performed in a single cycle followed by visible staining explained as described above.

| Antibody target protein | Vendor | Catalog no. | Clone | Fluorophore | Research resource identifier | Dilution |
| --- | --- | --- | --- | --- | --- | --- |
| **α-SMA** | Abcam | ab184675 | 1A4 | Alexa Fluor® 488 | AB_2832195 | 1:50 |
| **CD3E** | NovusBio | NBP2-54392AF647 | C3e/1308 | Alexa Fluor® 647 | AB_3095415 | 1:50 |
| **CD8A** | Thermo Fisher Scientific | 53-0008-80 | AMC908 | Alexa Fluor® 488 | AB_2574412 | 1:50 |
| **CD20** | eBioscience | 53-0202-80 | L26 | Alexa Fluor® 488 | AB_10734357 | 1:50 |
| **CD68** | CST | 79594 | D4B9C | PE | AB_2799935 | 1:50 |
| **E-cadherin** | CST | 3199 | 24E10 | Alexa Fluor® 488 | AB_10691457 | 1:50 |
| **Pan Cytokeratin** | Thermo Fisher Scientific | 41-9003-80 | AE1/AE3 | eFluor™ 570 | AB_11218704 | 1:50 |
| **Vimentin** | NovusBio | NBP1-97670AF750 | V9 | Alexa Fluor® 750 | AB_10013743 | 1:50 |
| **COL1A1** | NovusBio | NB600-408AF750 | Polyclonal | Alexa Fluor® 750 | AB_10000511 | 1:50 |
| **Ki67** | CST | 11882 | D3B5 | Alexa Fluor® 488 | AB_2687824 | 1:50 |
| **CD163** | Miltenyi | 130-127-908 | REA1309 | PE | AB_2928274 | 1:50 |
| **E-cadherin** | Cell Signalling | 3195 | 24E10 | - | AB_2291471 | 1:200 |
| **HLA-G** | EXBIO | 11-291-C100 | MEM-G/1 | - | AB_10734353 | 1:200 |
| **Anti-goat Ms IgG** | Thermo Fisher Scientific | A21235 | - | Alexa Fluor® 647 | AB_2535804 | 1:1000 |
| **Anti-goat Rb IgG** | Thermo Fisher Scientific | A11008 | - | Alexa Fluor® 488 | AB_143165 | 1:1000 |

### Image analysis and contour export for laser microdissection

#### Lung cancer

Immunofluorescent and H&E staining were overlayed using the Warpy extension (v0.2.6) in Qupath (v0.4). ROI selection was based on panCK, Vimentin and E-Cadherin staining. Annotations were exported as geoJSON files and converted to the LMD compatible format (.xml) using the QuPath-to-LMD webapp.

#### Triple negative breast cancer

Triple-negative breast cancer whole slide images were processed with MCMICRO: Tiles were illumination corrected with BASIC v1.1.1, then stitched and registered with ASHLAR v1.17.1. Channels were background-subtracted from autofluorescence images using Backsub v0.4.1. The image stack was processed using a modified SOPA snakemake pipeline. The nuclear signal from the first cycle was segmented using cellpose v2’s “nuclei” model, with parameters: diameter 25, flow_threshold 0.8, cellprob_threshold –8, min_area 250, clip_limit 0.2, gaussian_sigma 1. For whole-slide images, we used SOPA’s patchify function to tile images into 5000 pixels tall/wide squares with 100 pixel overlap to distribute segmentation tasks across HPC. Segmentation masks were expanded by five pixels and used to quantify mean, standard deviation, and three quantiles (25% 50% 75%) of marker signal per cell. For quality control, image stacks were scaled to 8-bit and pyramidized with tile-size 4096. The following additional filtering steps were applied with visual quality control: (1) removal of manually labelled artifact regions, (2) removal of cells with too low/high nuclear signal, (3) removal of cells with too small/large areas, (4) removal of dropout cells where DAPI ratio between first and last cycle was not between 0.15-1.05. Cell phenotyping was done using the napari-cell-gater (github.com/CosciaLab/napari-cell-gater) to determine marker thresholds and scale marker signals for binomial distribution. The plugin allows the user to load an image, segmentation mask, and quantification matrix of any sample(s); then to recursively determine a threshold for each marker-sample combination. This is done with dynamic feedback, and by choosing any two features (intensity or morphology based) to populate a scatterplot. Lastly, we used scimap’s phenotype function using the generated signature-matrix after marker thresholding to label the cells. Artifacts present in the CD8 channel were manually removed prior analysis. For cellular neighborhood analysis, AnnData objects were merged and analyzed by scimap’s spatial_lda function with knn=30 and k=7.

### Laser microdissection and proteomic sample preparation

#### General procedure

We used the Leica LMD7 system with software version 8.3.0.08259. Tissue was collected using the 20x objective (HC PL FL L 20x/0.40 CORR) for lung cancer and breast cancer samples, and 63x objective (HC PL FLUOTAR L 63x/0,70 CORR PH2) for placenta and scDVP samples. Laser settings were optimized per tissue type and listed below. Contours were collected into low-binding 384-well plates (Eppendorf 0030129547) or into proteoChip EVO96 (C-PEVO-96-10). To ensure tissue settling at the bottom of the wells, 20μL of acetonitrile was added, followed by centrifugation for 2min at 2000g and vacuum drying for 10 min at 60 °C. Wells were finally inspected in the LMD7 system to confirm collection.

Sample purification was performed using Evotips Pure (Evosep) following manufacturer’s instructions, and samples were kept with Buffer A until LC-MS measurement.

#### First trimester placenta

Contours were collected by laser microdissection using the 63x magnification objective. About 150 cells were collected per sample, in total about 50,000 µm^2^. Cell lysis was performed with 4 μL of 60 mM TEAB, followed by brief centrifugation (2,000 RCF, 1 min) and heating at 95 °C for 60 min in a Bio-Rad thermal cycler (384-well module, lid at 110 °C). ACN (1 μL; 20% final) was added, and the samples were incubated at 75 °C for 60 min. After cooling, 2 μL LysC (2 ng/μL in LC-MS grade H_2_O) was added and digestion was performed at 37 °C for 4 h. Trypsin (2 μL, 2 ng/μL; Promega Trypsin Gold) was added for overnight digestion at 37 °C. Digestion was stopped with 1% TFA (v/v), and samples were vacuum-dried (∼60 min, 60 °C) and stored at –20 °C. Before LC-MS, peptides were purified using Evotips Pure, eluted into a 96-well plate, vacuum-dried, and resuspended in 4.2 μL MS loading buffer (3% ACN, 0.2% TFA); the plate was vortexed (10 s), centrifuged (5 min, 2,000 RCF) and 4μL injected into the EASY-nLC-1200 system.

#### Lung cancer

Contours were collected by laser microdissection using the 20x magnification objective. All samples’ total area ranged between 50,000-100,000 µm^2^. Cell lysis and protein digestion was performed as described above for the placenta samples. After trypsin digestion, samples were vacuum-dried (∼60 min, 60 °C) and stored at –20 °C. Immediately before measurement, C18 based peptide purification (Evotip Pure) was performed, and samples were maintained in Buffer A prior LC-MS analysis.

#### Triple negative breast cancer

The total area for each sample was 25,000 µm^2^. Protein extraction and digestion used a DDM-based protocol as described above. A lysis buffer containing 0.025% DDM, 5 mM TCEP, 20 mM CAA, and 0.1 M TEAB in water was dispensed into wells (4 μL per sample) using a MANTIS Liquid Dispenser. Plates were sealed with PCR ComfortLid seals and incubated at 95 °C for 60 min. After cooling, 2μL of LysC (4 ng/μL in 0.1 M TEAB [pH 8.5] with 30% acetonitrile) was added, and samples were digested for 4 h at 37 °C in a thermal cycler. Subsequently, 3μL of trypsin (2ng/μL in 0.1 M TEAB [pH 8.5] with 10% acetonitrile) was added, and digestion proceeded overnight at 37 °C. Samples were vacuum dried for peptide purification.

### Liquid chromatography and mass spectrometry (LC-MS)

#### First trimester placenta

Peptides were analyzed using an EASY-nLC 1200 (Thermo Fisher) coupled to a trapped ion mobility spectrometry quadrupole time-of-flight mass spectrometer (timsTOF SCP, Bruker Daltonik). Samples were loaded onto a 20 cm in-house packed C18 column (75 μm ID, 1.9 μm ReproSilPur C18-AQ silica beads, Dr. Maisch, Germany) and separated at 250 nL/min with a gradient of increasing concentrations of buffer B (0.1% formic acid, 90% ACN in LC-MS grade H2O) to 60% buffer A (3% ACN, 0.1% formic acid in LC-MS grade H2O). The gradient duration was 21 min. MS acquisition was performed in diaPASEF mode using the vendor’s default ‘3×8’ method. Eight dia-PASEF scans were conducted, each divided into three ion mobility windows, spanning a mass-to-charge ratio (m/z) range of 400-1000 with 25 Th windows, and covering an ion mobility range from 0.64 to 1.37 Vs cm-2. The mass spectrometer was set to high sensitivity mode, featuring an accumulation and ramp time of 100 ms, a capillary voltage of 1750V, and a collision energy that increased linearly from 20 eV at 1/K0 = 0.6 Vs cm-2 to 59 eV at 1/K0 = 1.6 Vs cm-2. The collision energy was adjusted linearly based on ion mobility, starting at 59 eV at 1/K0 = 1.6 V s cm-2 and decreasing to 20 eV at 1/K0 = 0.6 V s cm-2.

#### Lung cancer

Lung cancer samples were directly eluted from Evotips using the Evosep One coupled to a trapped ion mobility spectrometry quadrupole time-of-flight mass spectrometer (timsTOF Ultra, Bruker Daltonik). Separation was performed using an Evosep 30SPD gradient with an IonOpticks Aurora Elite column. Samples were measured in dia-PASEF acquisition mode. The MS acquisition settings were the same as described above.

#### Triple negative breast cancer

Liquid chromatography was performed using the Evosep One LC system connected to a trapped ion mobility spectrometer with a quadrupole time-of-flight mass spectrometer (timsTOF Ultra 2, Bruker Daltonik). Separation was performed using an Evosep 80SPD gradient with an IonOpticks Aurora Rapid column. Samples were measured in dia-PASEF acquisition mode in high sensitivity mode with the MS method details as described above.

### Proteomic raw file and data analysis

#### First trimester placenta

Raw data were processed in DIA-NN (v1.8.1) using a predicted human spectral library (UniProt, 2021). Default settings were used with adjustments: mass range 100–1,700 m/z, precursor charge states 2–4, max 2 miscleavages, MS1/MS2 mass accuracy 15 ppm, match-between-runs enabled, and quantification set to ‘Robust LC’. Protein-level outputs (pg_matrix.tsv and unique_genes_matrix.tsv) were analysed in Perseus ^68^(v.1.16.0.5). Missing values were imputed from a normal distribution (width = 0.3, downshift = 1.8) after filtering for ≥70% quantified values per cell type group. Matrices were further analyzed in R (v4.1.2). To identify phenotype-specific protein markers, highly variable proteins were selected based on an ANOVA F-values greater than four. Differential abundance was assessed using Welch’s two-sided t-test followed by Benjamini–Hochberg FDR correction. Identified marker proteins (FDR < 0.05) overlapping with public transcriptomic datasets were visualized using DiVenn ^69^. Receptor ligand communication analysis was done using CellChat and included receptors and ligands indexed as “secreted,” “cell-cell contact” or “ECM-receptor” in their database. We used standard parameters (excluding per-protein smoothing) of 1000 permutations for the probabilistic inference of each cell pair and transmitter-receiver interaction, considering heterometric structures and interactor mediator proteins.

#### Lung cancer

Raw files were analyzed in DIA-NN (version 1.8.1), using a predicted human spectral library (UniProt, 2021) with small changes to default parameters. Briefly, mass range 400-1000, missing cleavages 2, max variable modifications 1, Ox(M) and Ac(N-term) modifications, precursor charge range 2-4, precursor m/z range 400-1000, fragment ion m/z range 200-1000, mass accuracy 15, MS1 accuracy 15, and MBR active. Raw files were pre-processed in Perseus as mentioned above. Missing values were imputed from a normal distribution (width = 0.3, downshift = 1.8) after filtering for ≥70% quantified values per cell type group. Downstream analysis was performed in R (v4.3.2). Differential abundance was assessed using Welch’s two-sided t-test followed by Benjamini–Hochberg FDR correction of 5%. For comparisons with RNA levels, the spatial transcriptomics dataset previously published in collaboration with Pentimalli et al. ^37^ was used. For analyses in figures 3i-k, publicly available data from the LUAD TCGA dataset was used.

#### Triple negative breast cancer

Raw files were analyzed in DIA-NN (version 1.9) using a predicted human spectral library (UniProt, release 2022) with small changes to default parameters. Briefly, maximum number of variable modifications 1, mass accuracy 15, MS1 accuracy 15, and MBR active.

Proteomic data analysis was performed in Python v3.12.8, using AnnData v0.11.2 and SpatialData v0.4.0 as the main data formats. Analysis was performed using opendvp functions with geopandas v1.0.1 and pandas v2.2.3 in the background. Plotting was performed using matplotlib v3.10.0, seaborn v0.13.2, and pycomplexheatmap. The DIA-NN output was processed and adapted to the AnnData object using the openDVP DIANN_to_adata function, and quality control samples were filtered out. Proteins that did not have at least 70% valid values in any cellular neighborhood were filtered out. The remaining NaN values were imputed by low-abundance normal distribution per protein (width=0.3, downshift=1.8). Scanpy v1.11.0 was used to perform PCA dimensionality reduction and plotting. Hierarchical clustering and plotting of heatmap took place with pycomplexheatmap v1.8.1, using method “average,” and metric “cityblock.” T-tests and ANOVA were performed using pingouin v0.5.5. Spearman’s correlation was calculated using scipy v1.16.

## Data availability

Mass spectrometry proteomics data have been deposited to the ProteomeXchange Consortium via the PRIDE partner repository ^70^ with the identifier PXD072882. RNA-seq data were obtained from GEPIA ^41^, an interactive web server for analyzing RNA sequencing expression data from the TCGA and GTEx projects, using a standard processing pipeline. The LUAD RNA-Seq dataset GEPIA used was based on the UCSC Xena project (http://xena.ucsc.edu).

Image data have been deposited to the BioImage Archive repository with the accession number S-BIAD-2194.

## Code availability

The source code of openDVP is fully opened and accessible on GitHub, including example tutorials: https://github.com/CosciaLab/openDVP. Extensive API documentation of functions can be found at: https://coscialab.github.io/openDVP/index.html. The QuPath-to-LMD webapp code can be found at https://github.com/CosciaLab/Qupath_to_LMD. Lastly, napari-cell-gater can be found at https://github.com/CosciaLab/napari-cell-gater.

## Declaration of generative AI and AI-assisted technologies in the writing process

During the preparation of this manuscript, the authors used Paperpal to improve the readability and language of the text. After using this tool, the authors reviewed and edited the content as needed and take full responsibility for the content of the publication

## Extended Data Figures

**Extended Data Fig. 1.**
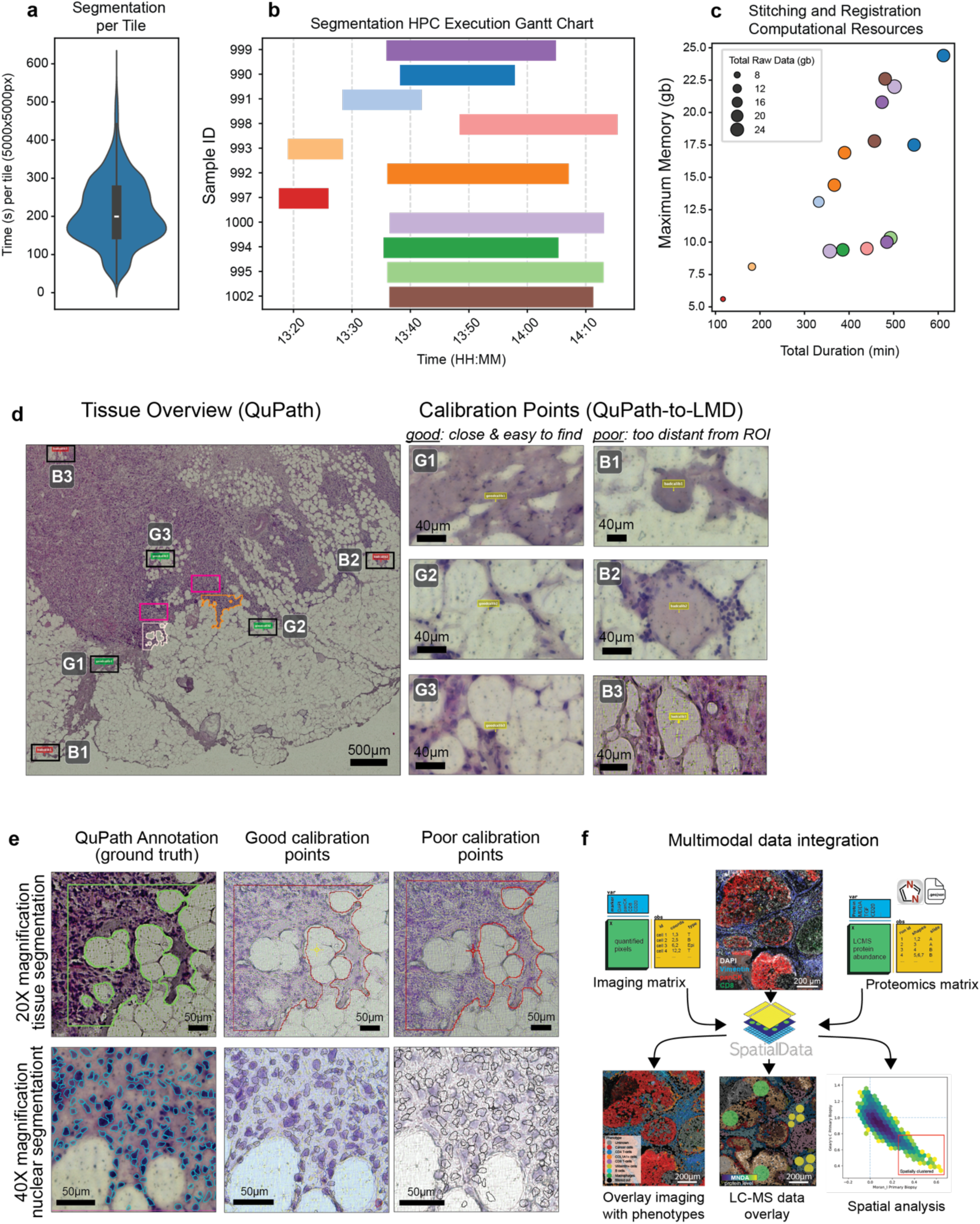
Benchmarking of computational resources and accurate reference point selection for reliable contour transfer for laser microdissection. **(a)** Violinplot showing duration in seconds of tile (5000×5000px) segmentation with Cellpose2 “nuclei” model. The central line indicates the median, while the box represents the interquartile range (IQR, 25th to 75th percentiles), whiskers extend to 1.5* IQR of the box **(b)** Gantt chart showing parallelization capabilities of MCMICRO and SOPA, 11 whole-slides with a total of 6+ million cells were segmented in less than two hours. **(c)** Bubble scatterplot with proportional computational resources to input dataset size. Stitching and registration via ASHLAR, is the slowest process in image processing pipeline. **(d)** Tissue overview and reference point definition in QuPath. G1–G3 indicate three good calibration points, located close to the objects of interest and easily identifiable. B1–B3 indicate three poor calibration points, which are located farther away and are difficult to recognize. The center shows four annotated regions: two magenta boxes corresponding to regions subjected to nuclear (H&E-based) segmentation in QuPath, and two irregular regions corresponding to tissue segmentation annotations. **(e)** Accurate annotation transfer depends on reliable calibration points. Top row: large tissue segmentation annotation and its contour transfer to the LMD7. Poor calibration points result in an upward shift of the entire contour. Bottom row: nuclear segmentation and contour transfer. In this case, poor calibration points lead to relatively large positional errors, which would result in improper single-cell collection. **(f)** Schematic illustrating how openDVP leverages the scverse SpatialData format to integrate antibody-based imaging, image-analysis–derived matrices, and LC–MS-based spatial proteomics data. This integration enables overlay of multiple information layers to guide laser microdissection and downstream spatial proteomics analyses.

**Extended Data Fig. 2.**
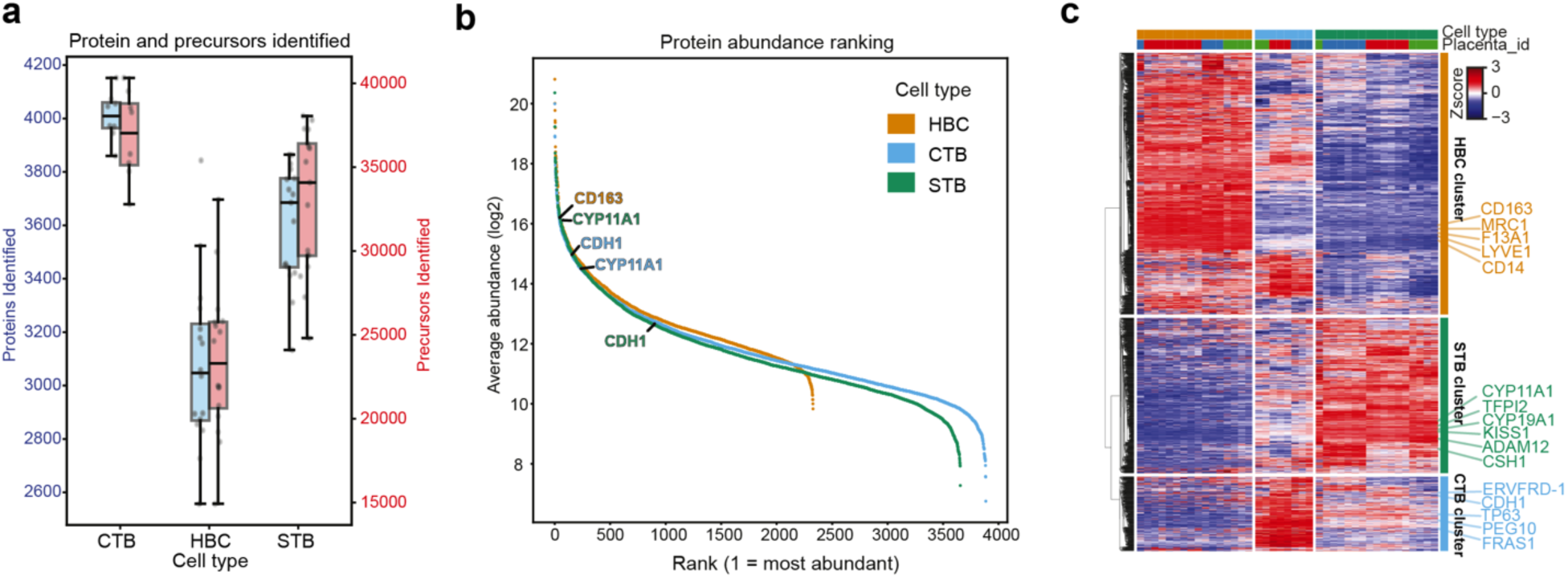
Proteome coverage and cell-type specific placental tissue proteomes. **(a)** Box plots showing the number of proteins (blue) and precursors (pink) identified per cell type; each dot represents a sample. The central lines indicate the median, the box the interquartile range, whiskers extend to (1.5*IQR) of the box. **(b)** Rankplot showing the dynamic range of proteins measured with labelled canonical markers and their rank for each cell type. **(c)** Unsupervised hierarchical clustering of 3,994 proteins, showing three distinct cell-type specific protein clusters used in Fig. 2h. Canonical markers of each cell type are labelled. The metric for clustering is “cityblock,” and the method is “average.” Clustering was performed by splitting the dendrogram into three largest groups. CTB, cytotrophoblast; STB, syncytiotrophoblast; HBC, Hofbauer cell.

**Extended Data Fig. 3.**
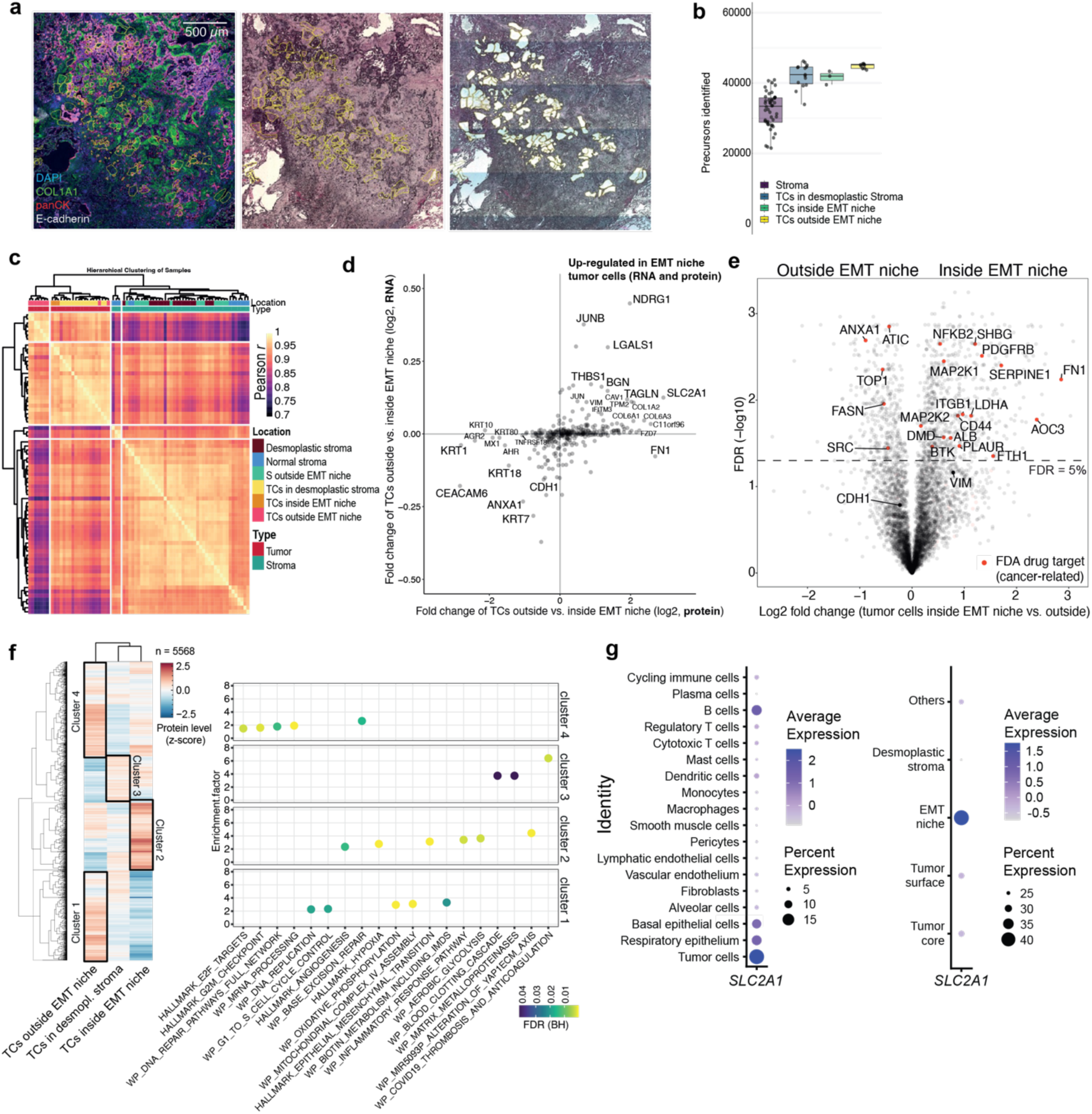
Integration of spatial transcriptomics and proteomics in lung cancer. **(a)** Imaging modalities (H&E and IF) used to guide laser microdissection. Yellow annotations show ROIs used for proteomic profiling. Left: Multiplex immunofluorescence used to locate tumor and stromal niches identified by spatial transcriptomics. H&E staining showing ROIs before (middle panel) and after LMD collection (right panel). **(b)** Boxplot showing the number of identified precursors per sample type. The central lines indicate the median, the box the interquartile range, whiskers extend to (1.5*IQR) of the box **(c)** Pearson’s correlation heatmap for all analyzed samples, method: “complete”, distance metric: “euclidean”. **(d)** Scatterplot comparing fold changes between tumor cells inside and outside the EMT niche, for both spatial proteomics and spatial transcriptomics. **(e)** Volcano plot showing differentially abundant proteins between tumor cells inside and outside the EMT niche as defined by CosMx spatial transcriptomics. FDA-approved drug targets are highlighted in red. E-Cadherin (*CDH1*) and Vimentin (*VIM*) are shown for reference. **(f)** Heatmap showing clusters of proteins between tumor cells inside the EMT niche, outside the EMT niche, and in the desmoplastic stroma, method: “complete”, distance metric: “euclidean”. The dot plot shows enriched Hallmarks and WikiPathways pathways for the different clusters using ClusterProfiler R package ^71^. **(g)** Dot plot of SLC2A1 expression across all cell types (left) and in tumor cells distributed in different niches (right). Dot size indicates the percentage of gene-expressing cells; color represents average expression (log-normalized and scaled).

**Extended Data Fig. 4.**
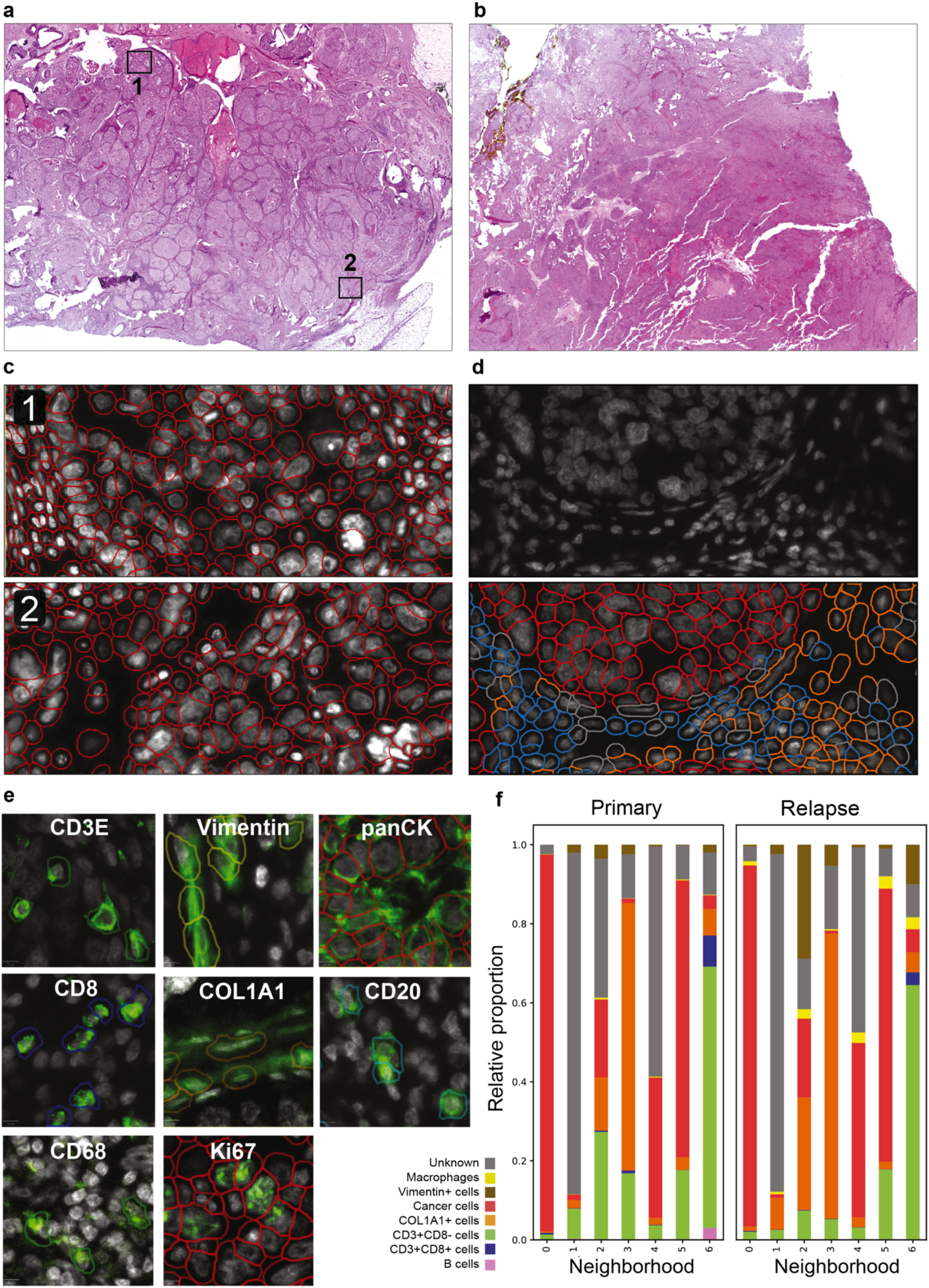
Consistent segmentation across large distances and staining variations. **(a)** H&E image of the primary TNBC tissue. **(b)** H&E image of the relapse TNBC tissue. Both tissues showed minor tissue losses due to cyclic imaging on PPS frame slides. **(c)** Nuclear stain overlayed with segmentation mask. These zoom-ins belong to distant regions from the primary TNBC tissue, areas shown and numbered in plot (a), showing segmentation consistency across entire whole-slide. **(d)** Zoom-ins of the primary tissue. The white color shows DAPI as the nuclear counterstain. Bottom image shows segmentation masks coloured by assigned phenotype labels. **(e)** Exemplary images for each IF marker used, overlaid with segmentation mask. Colors represent phenotyped cells for each marker (not for Ki67). **(f)** Barplots with cellular neighborhood compositions for each sample, showing the proportion of unclassified cells in grey. Related to Fig. 4g.

**Extended Data Fig. 5.**
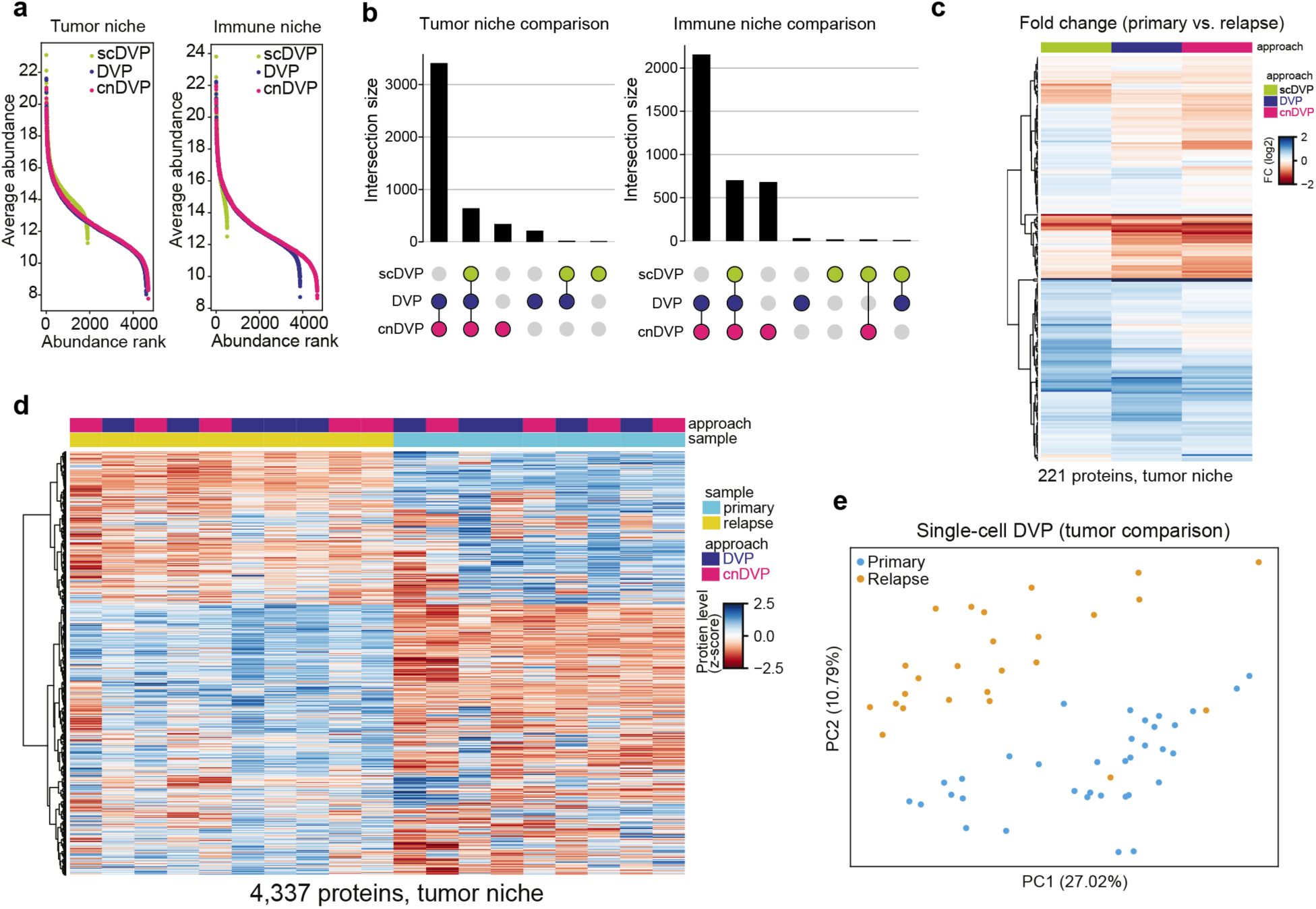
Comparative proteomic performance of complementary DVP strategies. **(a)** Rank plots illustrating the dynamic range of identified protein groups across three DVP approaches. Protein groups were retained if detected in at least five and three valid values for tumor and immune niches, respectively. **(b)** UpSet plots showing overlap of protein groups identified by the three DVP approaches. Only protein groups with at least 60% valid values per group are shown. **(c)** Heatmap of log₂ fold-change values between primary and relapse cancer samples for each DVP approach. Protein groups were stringently filtered to retain features with at least 70% valid values in each DVP approach. Missing values were imputed from a Gaussian distribution of lower-intensity values (mean shift −1.8, standard deviation 0.3, per feature). Hierarchical clustering was performed using Euclidean distance and the Ward linkage method. **(d)** Heatmap of log₂-transformed protein abundances in cancer samples profiled by the DVP and cnDVP approaches. Protein groups were retained if they contained at least 70% valid values in either the primary or relapse sample group. Missing values were imputed as in (c), and values were z-scored separately for the DVP and cnDVP approaches. Unsupervised hierarchical clustering was based on Euclidean distance and the Ward method. **(e)** Principal component analysis (PCA) of single-cell DVP (scDVP) cancer samples, colored by primary or relapse status. Protein groups were filtered to retain those with at least 50% valid values in either group (923 protein groups total). Missing values were imputed as described in (c).

**Extended Data Fig. 6.**
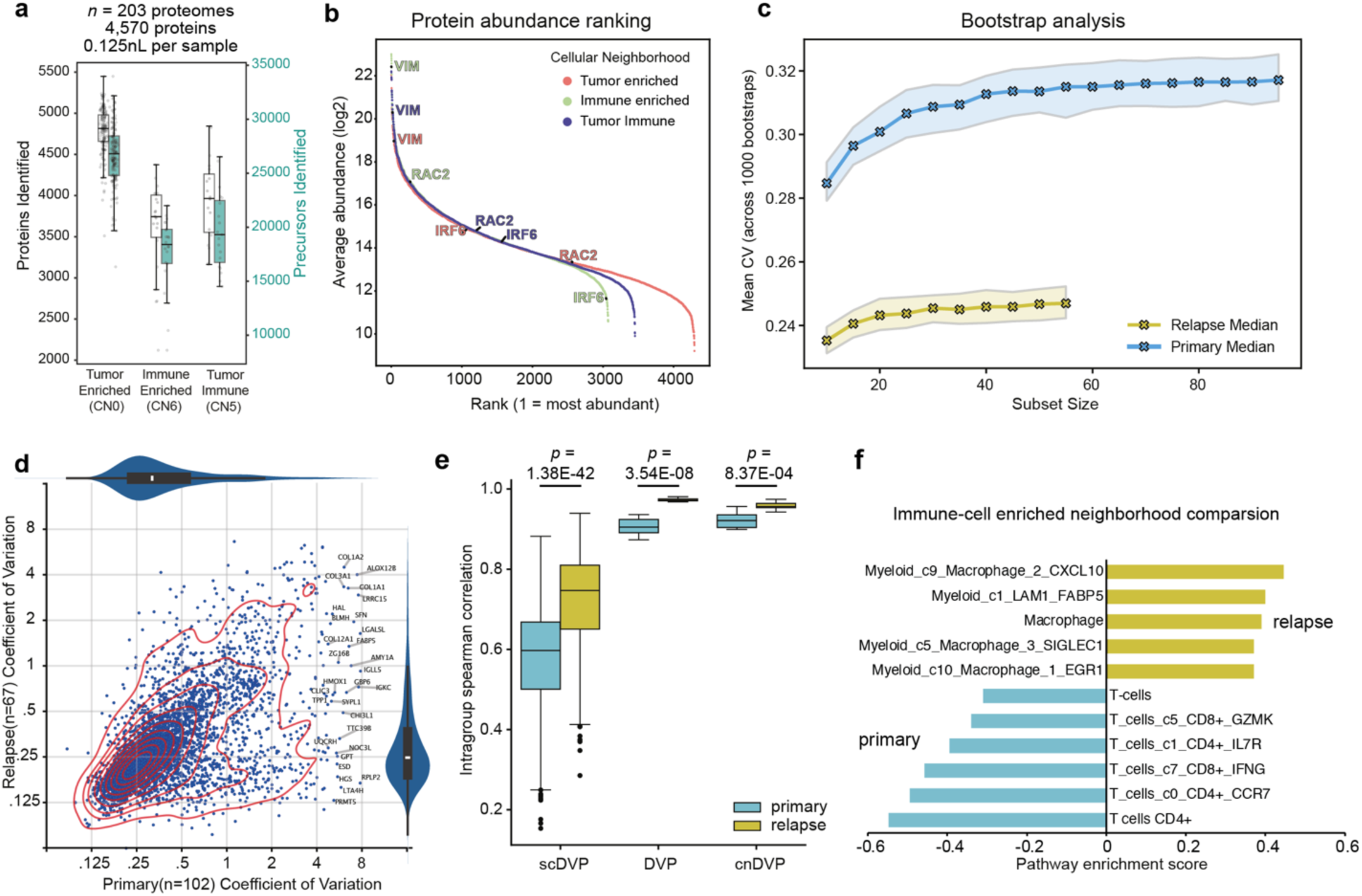
Neighborhood-guided DVP reveals proteomic differences between primary and recurrent TNBC. **(a)** Boxplot showing proteins and precursors identified per group of cellular neighborhoods. **(b)** Rankplot showing the dynamic range of protein abundance for each neighborhood. Vimentin (VIM) and RAC2 show the highest abundance in the immune-enriched samples, and IRF6 in tumor-enriched samples. **(c)** Line plot showing the spread of the mean CV of 1000 bootstrap simulations, repeated across different subset sizes for both samples. Note, plateauing was observed around 30-40 samples. **(d)** Scatterplot of intra-tissue protein level variability (CV) comparing the primary and relapse sample. Only samples of the tumor-specific CN0 were used analysis. Note, the primary tumor showed overall higher proteome variability. **(e)** Boxplots showing intragroup spearman correlation between cancer only samples for the three DVP approaches scDVP, DVP, and cnDVP. Protein groups are shown with at least 70% valid values for each sample group (primary or relapse) and approach (DVP, scDVP and cnDVP). P-values were calculated with a two-sided independent t-test. **(f)** Barplot of enriched cell type signatures between the primary and relapse sample. Only samples of the immune enriched CN6 were used for analysis. Breast cancer-specific cell type signatures were obtained from publicly available scRNAseq data described in Wu *et al*. ^56^. Note, the primary sample showed higher T cell signatures (e.g., CD4+, CD8+), whereas the relapsed immune niche featured higher myeloid/macrophage scores. For boxplots in (**a),(d)**, and **(e)** the central lines indicate the median, the box the interquartile range, whiskers extend to (1.5*IQR) of the box

